# The Gene Expression Deconvolution Interactive Tool (GEDIT): Accurate Cell Type Quantification from Gene Expression Data

**DOI:** 10.1101/728493

**Authors:** Brian B. Nadel, David Lopez, Dennis J. Montoya, Feiyang Ma, Hannah Waddel, Misha M. Khan, Serghei Mangul, Matteo Pellegrini

## Abstract

The cell type composition of heterogeneous tissue samples can be a critical variable in both clinical and laboratory settings. However, current experimental methods of cell type quantification (e.g. cell flow cytometry) are costly, time consuming, and can introduce bias. Computational approaches that infer cell type abundance from expression data offer an alternate solution. While these methods have gained popularity, most are limited to predicting hematopoietic cell types and do not produce accurate predictions for stromal cell types. Many of these methods are also limited to particular platforms, whether RNA-seq or specific microarrays. We present the Gene Expression Deconvolution Interactive Tool (GEDIT), a tool that overcomes these limitations, compares favorably with existing methods, and provides superior versatility. Using both simulated and experimental data, we extensively evaluate the performance of GEDIT and demonstrate that it returns robust results under a wide variety of conditions. These conditions include a variety of platforms (microarray and RNA-seq), tissue types (blood and stromal), and species (human and mouse). Finally, we provide reference data from eight sources spanning a wide variety of stromal and hematopoietic types in both human and mouse. This reference database allows the user to obtain estimates for a wide variety of tissue samples without having to provide their own data. GEDIT also accepts user submitted reference data, thus allowing the estimation of any cell type or subtype, provided that reference data is available.

**Author Summary:** The Gene Expression Deconvolution Interactive Tool (GEDIT) is a robust and accurate tool that uses gene expression data to estimate cell type abundances. Extensive testing on a variety of tissue types and technological platforms demonstrates that GEDIT provides greater versatility than other cell type deconvolution tools. GEDIT utilizes reference data describing the expression profile of purified cell types, and we provide in the software package a library of reference matrices from various sources. GEDIT is also flexible and allows the user to supply custom reference matrices. A GUI interface for GEDIT is available at http://webtools.mcdb.ucla.edu/, and source code and reference matrices are available at https://github.com/BNadel/GEDIT.

## Introduction

Cell type composition is an important variable in biological and medical research. In laboratory experiments, cell sample heterogeneity can act as a confounding variable. Observed changes in gene expression may result from changes in the abundance of underlying cell populations, rather than changes in expression of any particular cell type [1]. In clinical applications, the cell type composition of tissue biopsies can inform treatment. For example, in cancer, the number and type of infiltrating immune cells has been shown to correlate highly with prognosis ([2], [3], [4]). Moreover, patients with a large number of infiltrating T cells are more likely to respond positively to immunotherapy [5].

For many years, cell flow cytometry via FACS sorting has been the standard method of cell type quantification. More recently, single cell RNA-seq methods such as 10x Chromium, Drop-Seq, and Seq-Well have become available [6],[7]. However, both approaches suffer from significant limitations. FACS sorting is cumbersome and expensive, and some sample types require hours of highly skilled labor to generate data. Similarly, single cell RNA-seq methods remain expensive for large sample studies. Additionally, cell types such as neurons, myocytes, and adipocytes are difficult for these technologies to capture due to cell size and morphology.

Both FACS sorting and single cell methods have the potential to introduce bias, as these technologies require that tissue samples be dissociated into single cell suspensions. Many stromal cell types are tightly connected to one another in extracellular matrices. The procedures necessary to create single cell suspensions can damage some cells, while others remain in larger clusters that are not captured or sequenced. Consequently, subtle differences in sample preparation can produce dramatically different results [8,9]. While FACS sorting and single cell methods can produce pure samples of each cell type, the observed cell counts may not accurately represent the cell type abundances in the original sample. Tools like SCDC and MuSiC utilize single cell reference data to perform bulk deconvolution, but require that multi-subject single cell data be available for all the cell types of interest, which is not always the case [10,11].

During the past several years, digital means of cell type quantification, often referred to as cell type deconvolution or decomposition, have become a popular complement to FACS sorting and single cell approaches. However, these methods are still developing, and often suffer from limitations. For example, tools MCP-Counter and xCell allow for deconvolution of a set of predefined cell types, but do not support the inclusion of additional cell types or subtypes in a user friendly manner [12,13]. CIBERSORT is slow to run on large datasets, particularly if signature genes are not specified, and provides reference data only for hematopoietic cell types [14].

To overcome some of the limitations of existing cell abundance estimation tools, we present the Gene Expression Deconvolution Interactive Tool (GEDIT). GEDIT utilizes gene expression data to accurately predict cell type composition of tissue samples. We have assembled a library of reference data from 11 distinct sources and use these data to generate thousands of synthetic mixtures. In order to produce optimal results, these synthetic mixtures are used to test and refine the approaches and parameters used by GEDIT. We compare the performance of GEDIT relative to other tools using three sets of mixtures containing known cell type proportions: 12 *in vitro* mixtures of immune cells sequenced on microarrays, six RNA-seq samples collected from ovarian cancer ascites, and eight RNA-seq samples collected from blood. We also use GEDIT to deconvolute two sets of human tissue samples: 21 skin samples from patients with skin diseases, and 17,382 samples of varied tissues from the GTEx database. Lastly, we apply GEDIT to the Mouse Body Atlas, a collection of samples collected from various mouse tissues and cell types. We find that GEDIT compares favorably to other cell type deconvolution tools and is effective across a broad range of datasets and conditions.

## Results

### Reference Data

Reference data profiling the expression of purified cell types is a requirement for reference-based deconvolution. Methods that do not directly require reference data, such as non-negative matrix factorization, still require knowledge of expression profiles or marker genes in order to infer the identity of the predicted components. For this study, we have assembled or downloaded a set of 11 reference matrices, each containing the expression profiles of eight to 29 cell types (Table 1). These data sources span multiple platforms, including bulk RNA-seq, microarray, and single-cell RNA-seq. Complete details on the sources and assembly of these matrices are described in the methods [14–24].

**Table 1.**
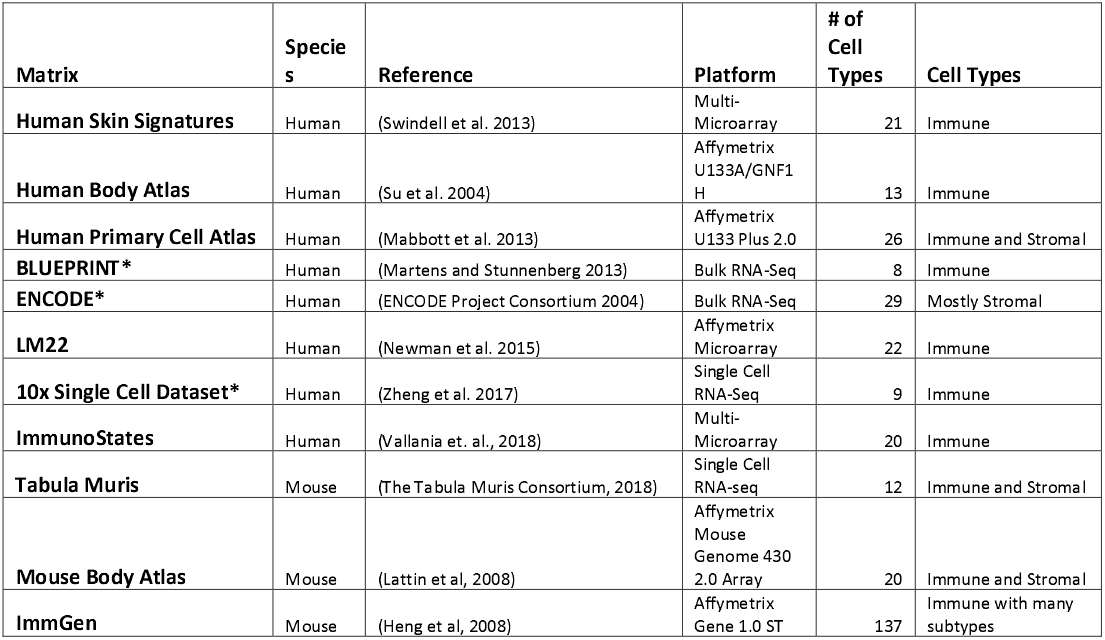
Library of Reference Data. Asterisk denotes matrices assembled from source data as part of this project. All matrices are compatible with GEDIT and available on the GitHub repository (https://github.com/BNadel/GEDIT).

### GEDIT Algorithm

GEDIT requires as input two matrices of expression values. The first is expression data is collected from the mixtures that will be deconvoluted; each column represents one mixture, and each row corresponds to a gene. The second matrix contains reference data, with each column representing a purified reference profile and each row corresponding to a gene. In a multi-step process, GEDIT utilizes the reference profiles to predict the cell type proportions of each submitted mixture (Figure 1).

**Figure 1.**
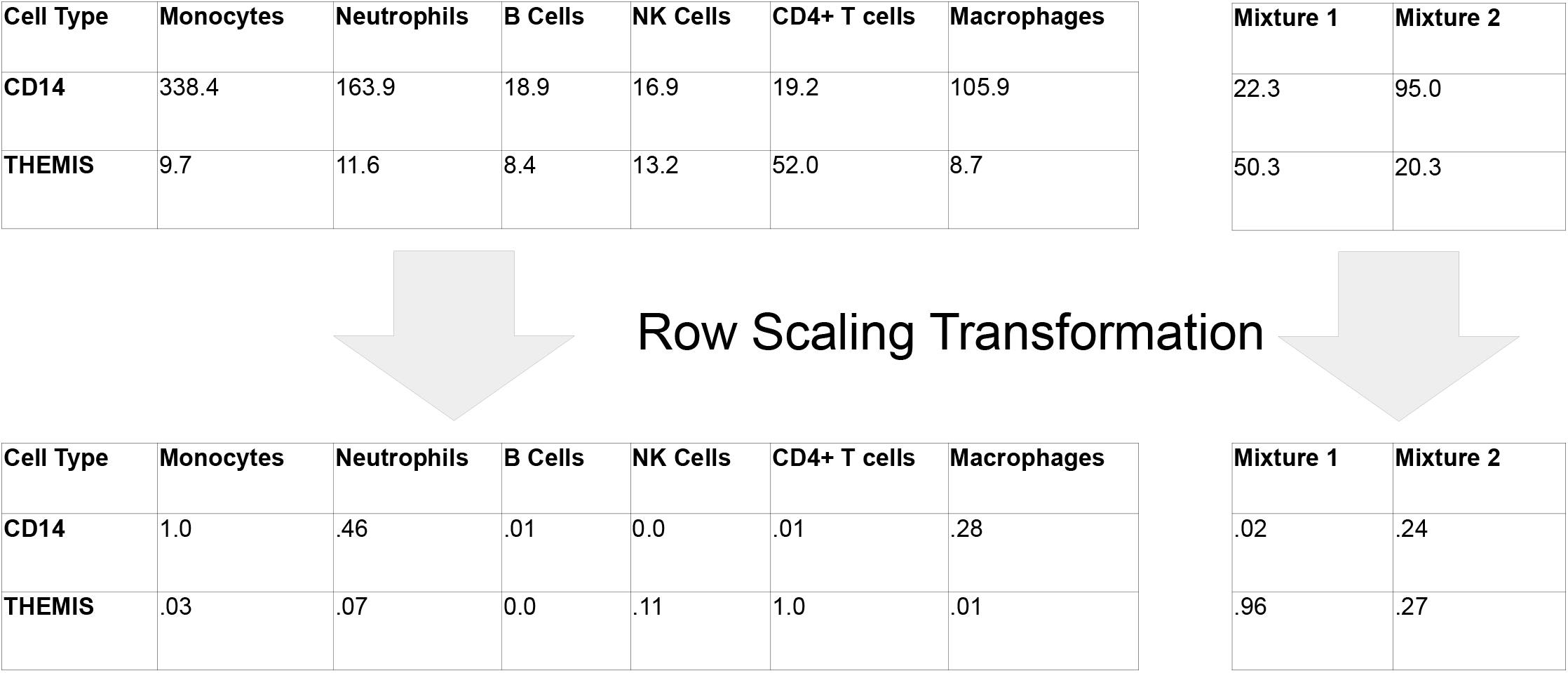
The GEDIT pipeline. The input matrices are quantile normalized then reduced to matrices containing only signature genes. Next, a row-scaling step serves to control for the dominating effect of highly expressed genes. Lastly, linear regression is performed, and predictions of cell type abundances are reported to the user.

### Parameter Tuning

We generated a large number of synthetic mixtures *in silico* to test the efficacy of GEDIT and to assess how accuracy varies as a function of four parameter choices (SigMeth, NumSigs, MinSigs, RowScale, described in Table 2). We produced a total of 10,000 simulated mixtures of known proportions using data from four reference matrices: BLUEPRINT, The Human Primary Cell Atlas, 10x Single Cell, and Skin Signatures. We then ran GEDIT on these simulated mixtures and evaluated its performance while varying four parameter settings (Figure 2) and other design choices. For this reason, these synthetic mixtures were not used to evaluate the performance of GEDIT relative to other tools. Instead, separate datasets were used for that purpose, as described in the section “Performance Comparison to Other Deconvolution Tools”. Based on these results, we selected default values for each parameter (SigMeth = Entropy, NumSigs = 50, MinSigs = 50, RowScale = 0.0). Full details on the generation of these simulations are described in the supplementary materials.

**Table 2.**
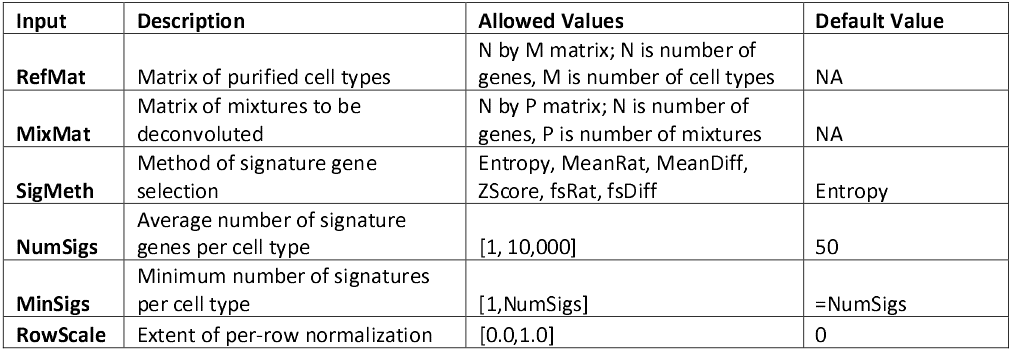
GEDIT inputs include two matrices and four parameter settings. RefMat is an expression matrix documenting the expression profiles of each cell type to be estimated. MixMat is an expression matrix documenting expression values for each sample to be deconvoluted. SigMeth determines the method by which signature genes are selected. NumSigs determines the total number of signature genes, whereas MinSigs sets the minimum number of signature genes for each cell type. RowScale refers to the extent to which expression vectors are transformed to lessen the dominating effect of highly expressed genes, with a value of 0.0 representing the most extreme transformation. Default values were determined by evaluating performance on a set of synthetic mixtures (Figure 2).

**Figure 2.**
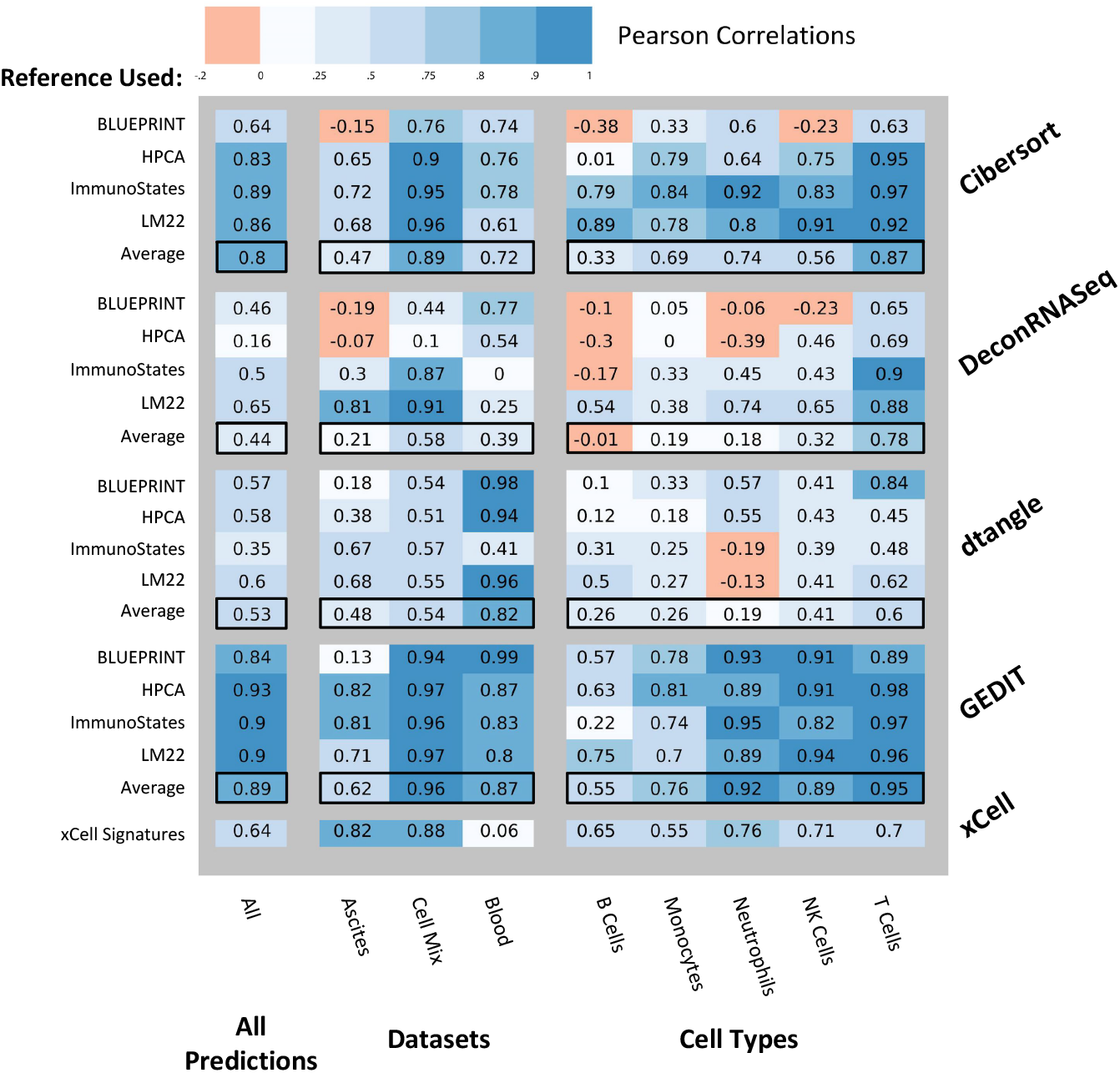
Effect of GEDIT parameter choices on accuracy of predictions in simulated experiments. 10,000 simulated mixtures were generated, each using one of four reference matrices, with either four, five, six, or ten cell types being simulated. Deconvolution was performed using a separate expression matrix than the one used to generate the mixtures. When not otherwise noted, we use the following parameters: signature selection method = entropy; number of signatures = 50; row scaling = 0.0; and number of fixed genes = number of signatures.

### Preprocessing and Quantile Normalization

The first step in the GEDIT pipeline is to render the two matrices comparable. This is done by first excluding all genes that are not shared between the two matrices. Genes that have no detected expression in any reference cell type are also excluded, as they contain no useful information for deconvolution. Both matrices are then quantile normalized, such that each column follows the same distribution as every other; this target distribution is the starting distribution of the entire reference matrix.

### Signature Gene Selection

GEDIT next identifies signature genes. Gene expression experiments can simultaneously measure tens of thousands of genes, but many of these genes are uninformative for deconvolution. Specifically, genes with similar expression levels across all cell types are of little use, as observed expression values in the mixtures offer no insight into cell frequencies. Genes that are highly expressed in a subset of cell types are more informative, and we refer to these as signature genes. By using only signature genes, rather than the entire expression matrix, the problem of deconvolution becomes more tractable and less computationally intensive. Moreover, identification of signature genes can be valuable to researchers for other applications (e.g. cell type assignment for scRNA-seq data).

In order to identify the best signature genes in a given reference matrix, GEDIT calculates a signature score for each gene. By default, this score is computed using the concept of information entropy. Information entropy quantifies the amount of information in a probability distribution, with highly uniform distributions having the highest entropy. The expression vector for each gene (i.e. the set of expression values across all cell types in the reference) is divided by its sum, such that the entries can be interpreted as probabilities. Information entropy is then calculated according to its mathematical definition (see Methods), and genes with the lowest entropy are selected as signature genes. Entropy is minimized when expression is detected only in a single cell type and maximized when expression values are equal across all cell types. Thus, by selecting genes with low entropy, we favor genes that are expressed in a cell type specific manner. By default, 50 signature genes are selected for each cell type in the reference matrix. We chose 50 signature genes, and entropy as our scoring method, because it returned optimal results when run on 10,000 synthetic mixtures (see Figure 2a,b).

We also evaluated the effect of accepting more signature genes for some cell types than others, depending on how many genes have low entropy. In this scheme, on average 50 signature genes are used per cell type. However, a fourth parameter is used, which specifies the minimum number of signature genes per cell type. After these have been selected, remaining signature genes are added based only on lowest entropy, regardless of cell type of maximal expression. We found that this parameter had minimal effect on accuracy, when applied to synthetic mixtures (Figure 2c). Therefore, this option is not used by default, though it can be specified by the user.

### Row Scaling and Linear Regression

One complication in the application of linear regression to gene expression data is the drastically different scale at which some genes are expressed. For example, CD14 and THEMIS (Figure 3) have both been identified as signature genes: CD14 for monocytes and THEMIS for CD4+ T cells. However, CD14 is expressed at much higher levels in most cell types and will have a larger impact on the estimation of cell type composition, relative to THEMIS. In other words, the possible penalty resulting from a poor fit of CD14 is much larger than the penalty from a poor fit of THEMIS.

**Figure 3.**
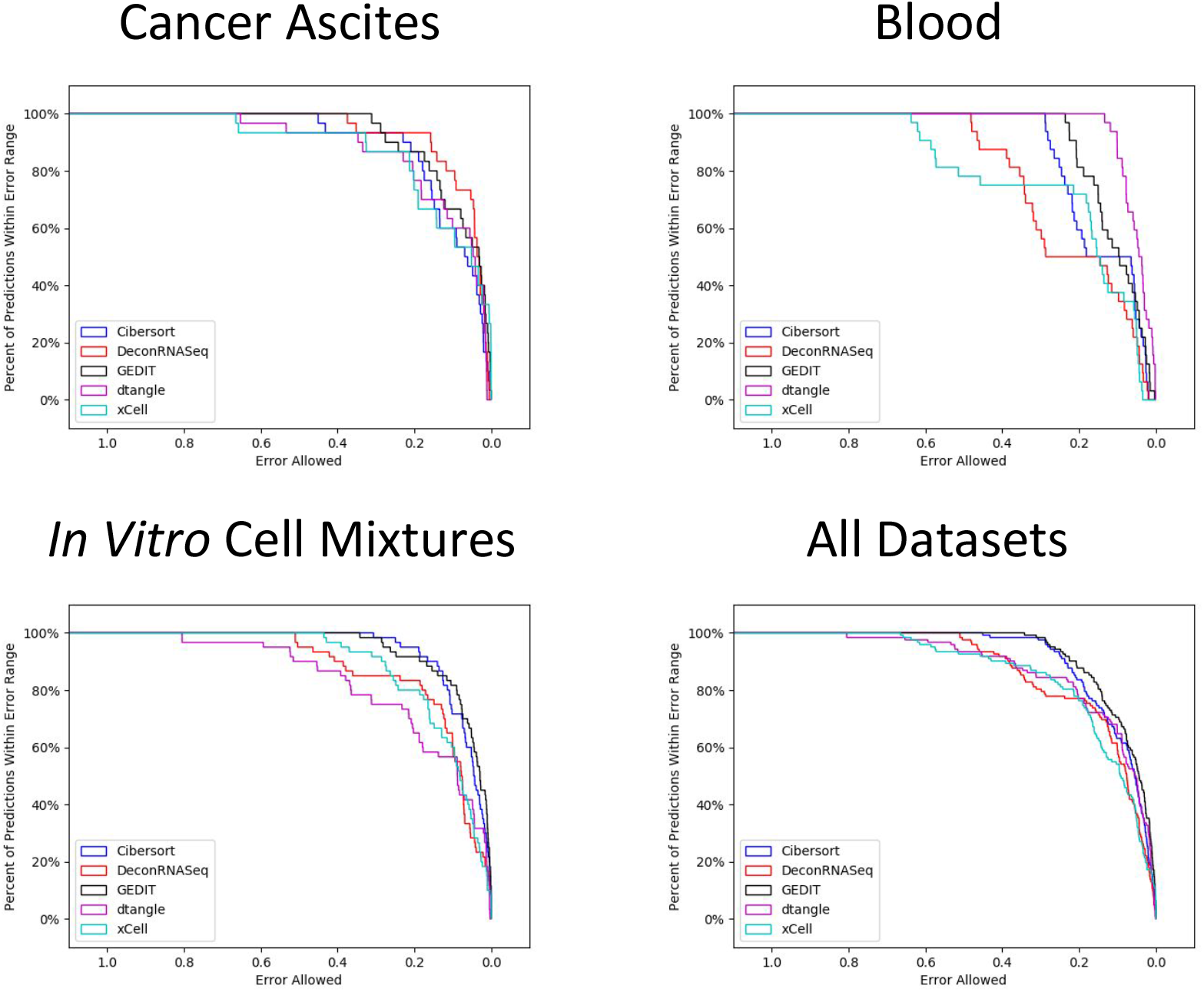
The “row scaling” transformation, as implemented by GEDIT. CD14 and THEMIS are two examples of signature genes with drastically different magnitudes of expression. CD14 is a signature gene for monocytes, and THEMIS for CD4+ T cells. The original expression vectors are transformed, such that all values fall between 0.0 and 1.0, equalizing the effect of genes with varying magnitudes of expression.

In order to equalize the effect of each signature gene on the linear regression, we implement a transformation that we term row scaling. Specifically, the range of all observed values for a particular gene (including reference cell types and samples) is adjusted such that the maximum value is 1.0 and the minimum value is 0.0. As a result, all genes have a comparable influence on the calculation of the linear regression solution, regardless of overall magnitude of expression. This transformation can be modulated by adjusting the row scaling parameter. By default, the value of this parameter is 0.0, and the transformation is applied as described above. Values between 0.0 and 1.0 are also allowed, which reduces the extent of the transformation (see Methods for details). Linear regression is then performed in R using the glmnet package, as described in the methods.

### Performance Comparison to Other Deconvolution Tools

In order to assess the performance of GEDIT relative to other tools, we perform an experiment comparing GEDIT to 4 other deconvolution tools on datasets of known cell-type content (CIBERSORT, DeconRNASeq, dtangle and xCell; [13,14,25,26]). Non-deconvolution tools like MCP-counter, SAVANT, and the DCQ algorithm are excluded from this study because they do not predict cell type fractions [12,27,28].Tools that require single cell data, such as MuSiC and CPM, are also excluded, as this study is limited to tools that operate on bulk expression data [11,29]. See Table 3 for a summary of current bulk deconvolution methods.

**Table 3.**
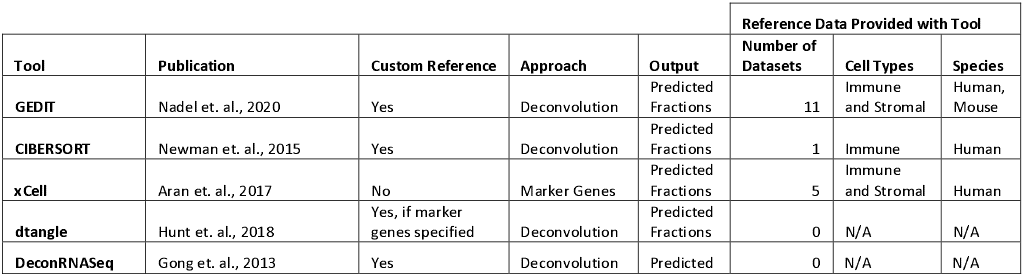
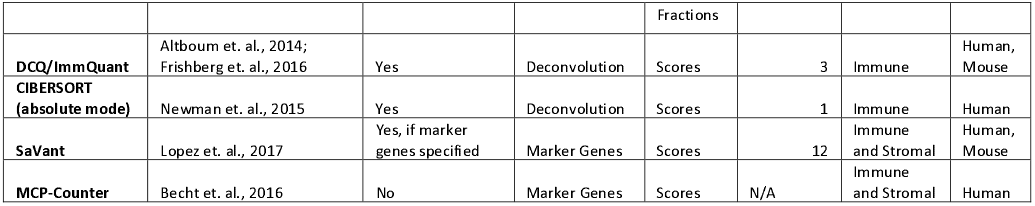
High level characteristics of current cell type estimation tools. Some tools accept custom references, which allows the tool to estimate the abundance of cell types not present in the default reference. Tools listed here take one of two approaches: they either perform deconvolution (most commonly regression) or calculate a score based on intensity of marker gene expression. Depending on the tool, the output can be interpreted as fractions corresponding to the abundance of each cell type, or as scores for each cell type that cannot necessarily be compared in an inter-cellular manner.

To perform this study, we utilize three datasets for which cell type fractions have been estimated using orthogonal methods. Two of these datasets were used in a recent benchmarking study [30]. Both are profiled using RNA-seq, and represent samples collected either from human cancer ascites or human blood [31,32]. In both cases, cell type fractions have been evaluated by FACS sorting. The final dataset was prepared *in vitro* and consists of six cell types that were physically mixed together (in known proportions) to prepare 12 mixtures. These mixtures were then profiled using an Illumina HT12 BeadChip microarray. Adding to the previous benchmarking study, we also explore the effect of using four separate reference datasets: The Human Primary Cell Atlas, LM22, ImmunoStates, and a reference constructed from BLUEPRINT data. For each dataset, all tools (except xCell) were run four times, each time using a different reference matrix. The optimal choice of reference matrix varies greatly depending on the exact combination of tool, dataset, and cell type. While using LM22 often produces the most accurate results, there are many exceptions. For instance, DeconRNASeq and GEDIT produce their best results for the blood dataset when using the BLUEPRINT reference. For the ascites data, several tools prefer ImmunoStates as the optimal reference choice. The best choice of reference is highly dependent on the nature of the input data and on the tool being used. In practice, researchers may wish to perform deconvolution multiple times--in each case using a separate reference matrix--and compare results for consistency.

Compared to the other tools, GEDIT produces robust and consistently accurate results (Figures 4,5). For many tools, the quality of predictions varies greatly depending on the cell type, dataset, or choice of reference matrix. When results are averaged across the four possible reference choices, GEDIT produces the minimum error and maximum correlation for all three datasets. This result suggests that GEDIT is a strong choice when researchers are using novel references matrices that have not been curated or tested.

**Figure 4.**
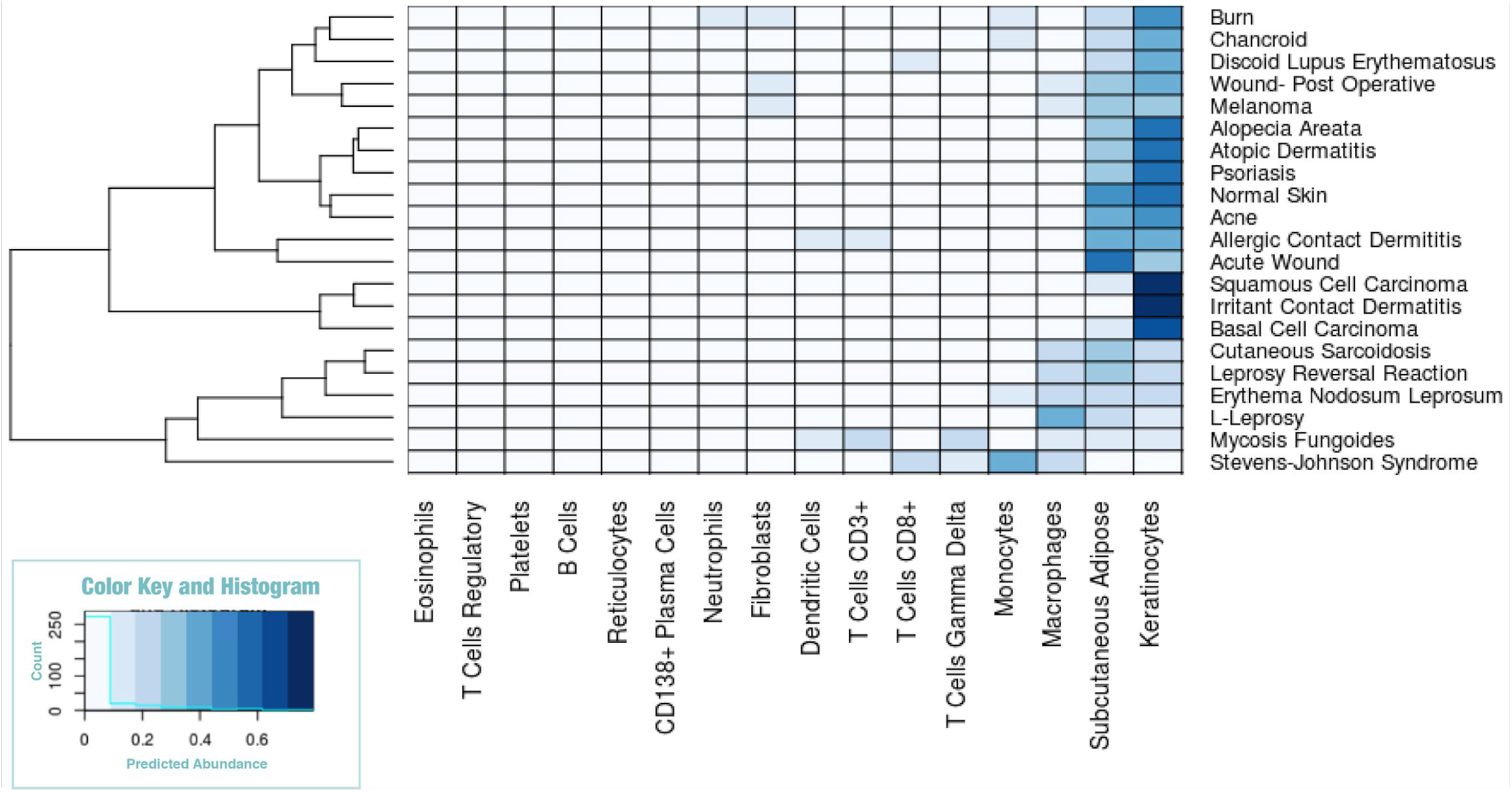
Performance of five deconvolution tools when applied to a set of 26 physical samples from three sources. Actual cell type fractions are either known due to controlled cell mixing (Cell Mix) or estimated by FACS sorting (Ascites and Blood). In each instance, we calculate the correlation between actual cell type fractions and those predicted by deconvolution; deeper blues represent higher correlations. We test four different reference datasets for each tool, and averaged correlations across these 5 cases are shown in boxes. We calculate correlations for each cell type (right 5 columns), for each of the 3 mixtures (middle 3 columns), and for all predictions regardless of cell type or data source.

**Figure 5.**
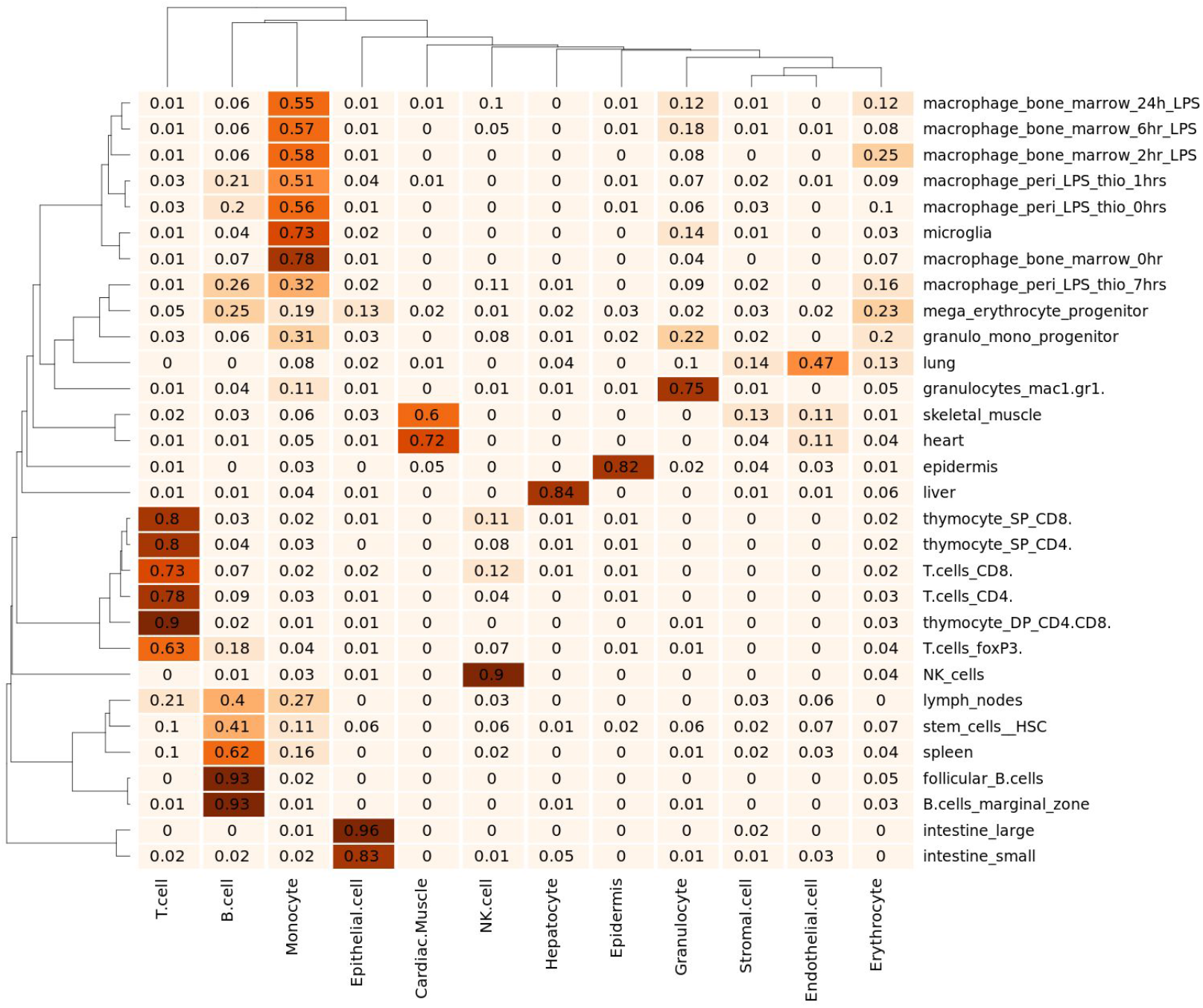
Distribution of errors, calculated as the difference between predicted and actual cell type fractions. Each point on the graph represents the percentage of predictions (y-axis) that are accurate within a particular error range (x-axis).

We also perform 2 additional comparisons between GEDIT and other deconvolution tools. Firstly, we create 100 simulated mixtures of pancreatic cells (alpha, beta, gamma, delta) using single cell data from a recent single cell experiment (details in supplementary materials). We evaluate the accuracy of each tool when used to predict the cell type content of these synthetic mixtures, and GEDIT provides the lowest overall error (Supplementary Figure 3).

Lastly, we perform an evaluation of runtime required for each tool. We randomly select batches of 100, 200, 500, 1000, and 2000 samples from the GTEx database, and measure CPU time required to deconvolute these batches for each tool. The runtime of GEDIT, dtangle, and DeconRNASeq scales well with growing input size, taking at most 20 minutes on average (Supplementary Figure 4).

### Skin Expression Data

We further validate GEDIT by using it to deconvolute a set of skin biopsies from humans with a variety of skin diseases [13]. The exact cell type composition of these samples is unknown, but we have reasonable expectations based on skin and disease biology. For example, macrophages are known to be abundant in granulomas of leprosy legions, and Steven-Johnson Syndrome produces blisters that fill with large numbers of monocytes [33,34]. We find that, in all cases, predictions made by GEDIT conform well with these biological expectations. Keratinocytes are highly predicted in most cases, as one would expect with skin samples (Figure 6). Deviations from this pattern correspond with disease biology. Monocytes are highly predicted in Stevens-Johnson syndrome, as are macrophages in the three leprosy samples, and T cells in the Mycosis Fungoides (T cell lymphoma) sample.

**Figure 6.**
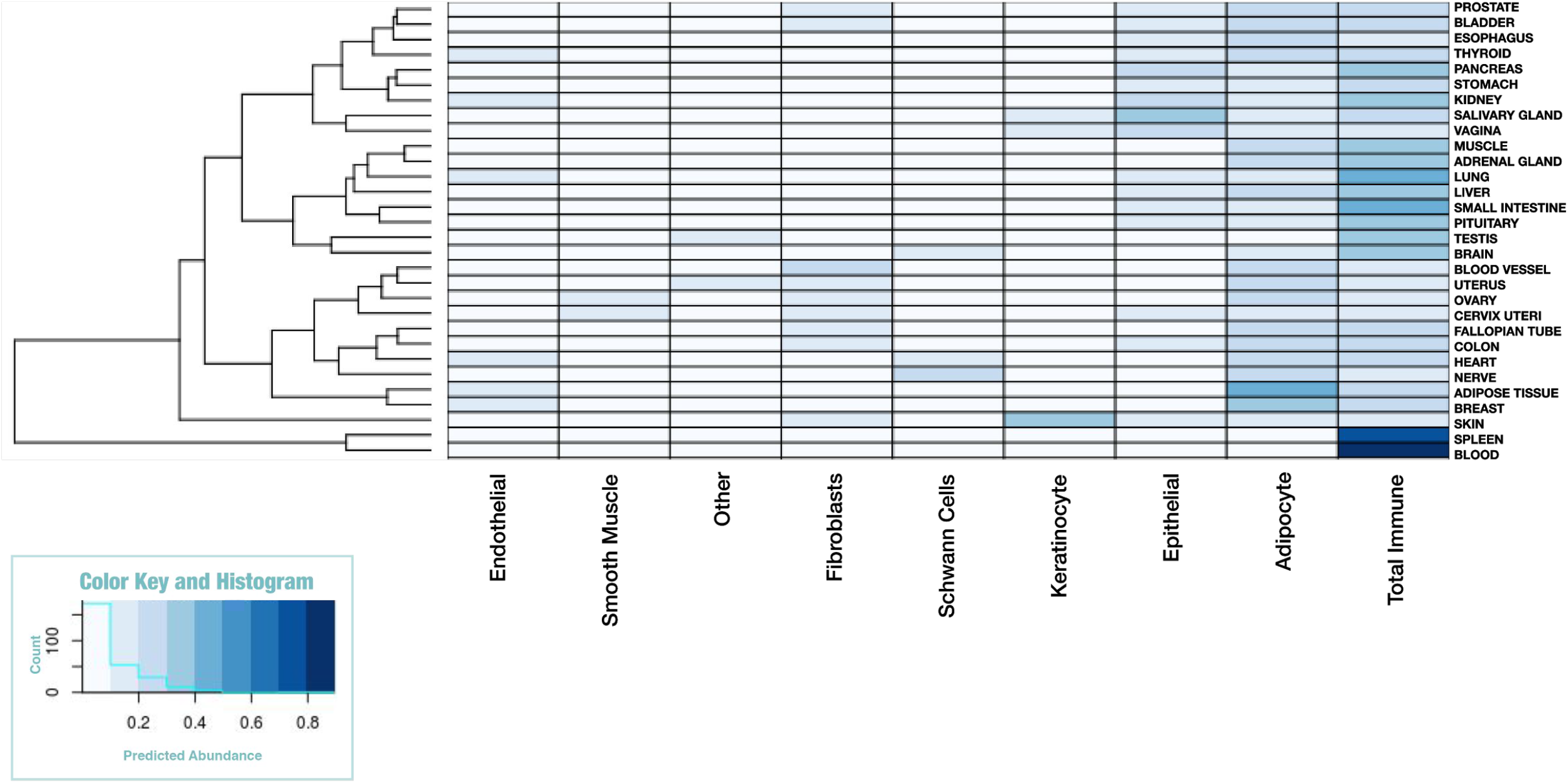
GEDIT predictions for 21 samples of various skin diseases. GEDIT correctly identifies keratinocytes and subcutaneous adipose as the most common cell. Deviations from this pattern correspond to disease biology. SJS represents blister fluid from Steven Johnson Syndrome, and is predominantly immune cells. LL and RR represent two forms of leprosy, which result in large numbers of macrophages. MF is a T Cell Lymphoma.

### Application of GEDIT to Mouse Data

GEDIT can be used to decompose data from any organism for which reference data is available. Here, we demonstrate the efficacy of GEDIT when applied to the Mouse Body Atlas, a collection of tissue and cell type samples collected from mice [23]. As reference data, we assembled a matrix of 12 cell types using single cell data from the Tabula Muris [20]. GEDIT correctly infers the identity of purified cell types, including six samples that consist of either pure NK cells, B cells, T cells, or granulocytes (Figure 7). An entry for macrophages is not available in the reference used, but most macrophage samples are identified as monocytes, which is the most similar cell type present in the reference matrix. For more complex tissues, GEDIT predicts cell type fractions that correspond to the biology of the samples. Hepatocytes are predicted to be highly prevalent in the liver sample (84%) and are not predicted in any other sample (less than 5% in all cases). Similar patterns hold for keratinocytes in the epidermis, epithelial cells in two intestinal samples and cardiac muscle cells in heart and muscle samples.

**Figure 7.**
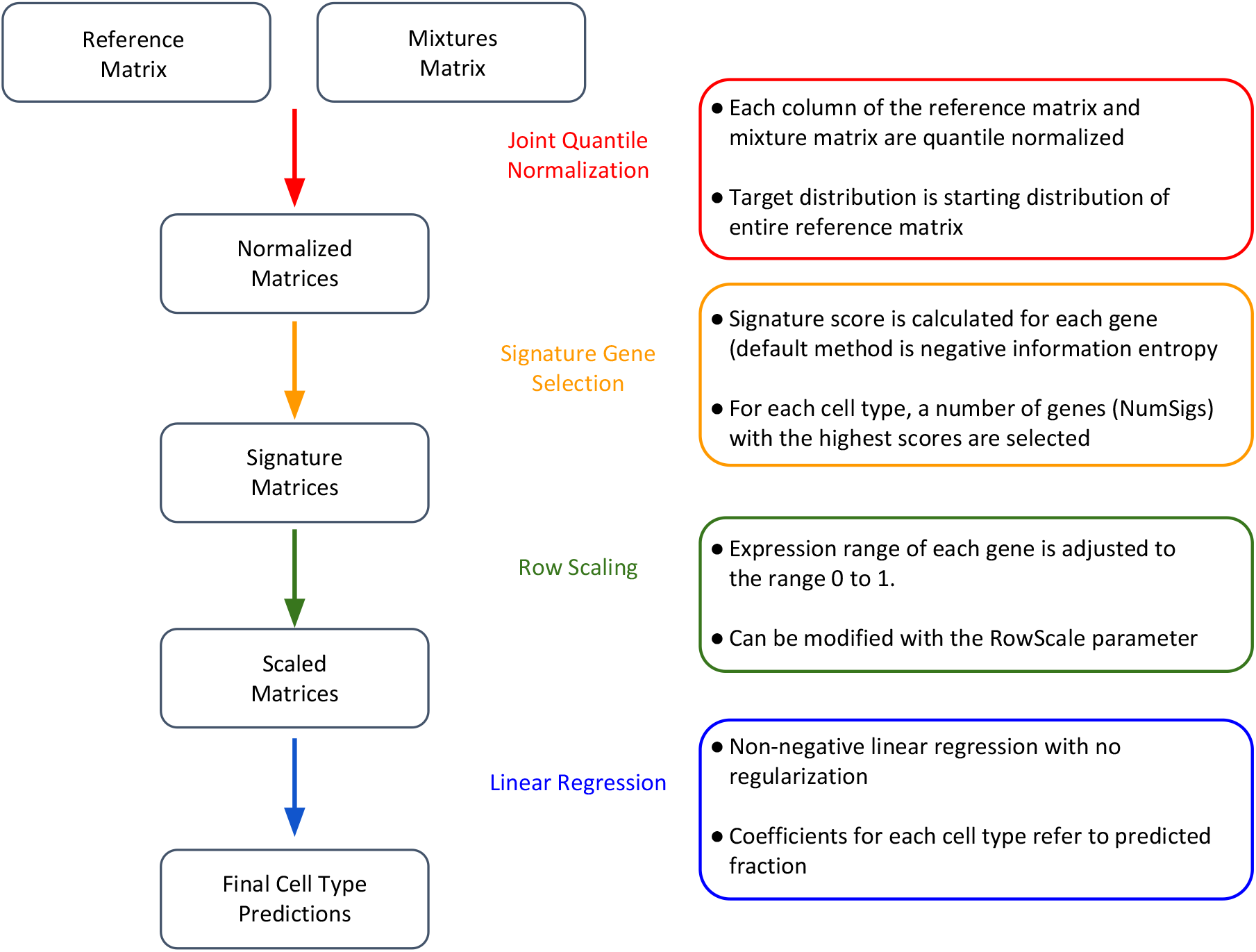
GEDIT predictions on 30 samples collected from various mouse tissues and cell types (mouse body atlas [23]). Predictions largely conform with tissue and cell biology.

### Deconvolution of GTEx Database

To assess the use of GEDIT across very large datasets, we applied the tool to 17,382 GTEx RNA-seq samples collected from various tissues. However, no single reference contained all relevant cell types. For example, none of the available references contain both myocytes and adipocytes (Supplementary Figure 1). Therefore, we predicted proportions three times using three separate references (BlueCode, Human Primary Cell Atlas, Skin Signatures). We then combined these outputs by taking their median value. This allowed us to produce predictions spanning a larger number of cell types than are present in any one reference matrix (Figure 8).

**Figure 8.**
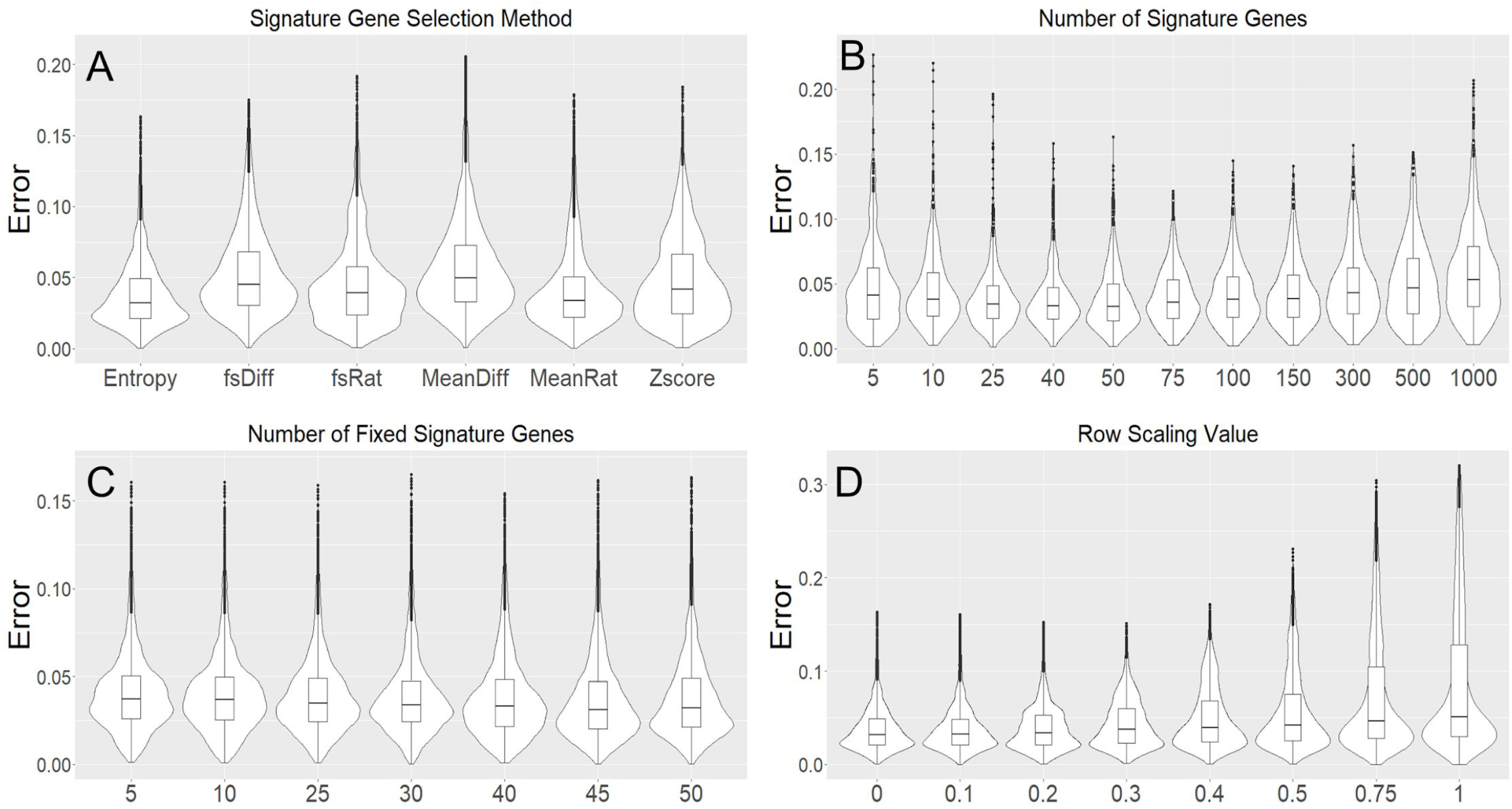
GEDIT cell type predictions when applied to 17,382 samples from the GTEx database. Here, predictions have been averaged for each tissue of origin.

These predictions largely conform to biological expectations. For example, immune cells are predicted to have high abundance in blood and spleen, adipocytes in adipose tissue, Shwann cells in nerve and heart, and keratinocytes in skin. Each of these patterns matches expectations of which cell types should be present in these tissues. Neither cardiac myocytes nor smooth muscle are highly abundant in GTEx muscle samples. This is likely because the GTEx samples are collected from skeletal muscle, which is known to have an expression profile that is distinct from that of cardiac and smooth muscle.

### GEDIT Availability

GEDIT can be run online at http://webtools.mcdb.ucla.edu/. Source code, associated data, and relevant files are available on GitHub at https://github.com/BNadel/GEDIT. We provide access to the tool, a set of varied reference data, and two sample mixture matrices. The website automatically produces a heatmap of predicted proportions for the user, as well as a .tsv file. The user also has access to the parameter choices of GEDIT (signature gene selection method, number of signature genes, row scaling).

## Methods

### GEDIT Algorithm

#### Signature Gene Selection

During signature gene selection, we automatically exclude genes with zero detected expression in half or more of cell types. Observed expression values of exactly zero are often the result of either technical artefacts or resolution issues. Using such genes as signatures can result in inaccurate and highly unstable results, particularly when working with scRNA-seq derived data. As an additional safeguard, we treat all remaining expression values of zero as the lowest observed non-zero value in the matrix. Implementing this change has minimal effect on most genes but prevents genes with resolution issues from achieving artificially high scores. We consider this transformation valid, since values of zero generally do not represent zero expression, but rather an expression level below the detection limit of the technology used.

For any given gene, a scoring method takes as input the vector of the expression values across all reference cell types, and outputs a score. A gene is considered a potential signature gene in cell type X if it is expressed more highly in X than any other cell type. For each cell type, we keep only the N genes with the highest scores, where N is the NumSigs parameter.

Information entropy (H) is calculated using the following formula:

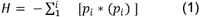

where *p*_i_ is the probability of the *i*^*th*^ observation. To apply this to expression values, we convert the vector of expression values into a vector of probabilities by dividing by its sum. In an equal mixture of each cell type, the *i*^*th*^ probability can be interpreted as the fraction of transcripts originating from the *i*^*t*h^ cell type.

#### Row Scaling

During this step, we apply a transformation on the expression values for each gene. Each gene has measured expression in N purified cell types and M samples. Each of these values, X_old_, is transformed according to the following formula:

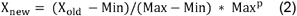

Where Min is the minimum of all M + N original values, Max is the maximum of those values, and p is a tunable parameter with natural range p ∈ [0.0,1.0]. This procedure produces values between the range of 0.0 and Max^p^.

#### Linear Regression

Non-negative linear regression was performed using the glmnet package in R. The glmnet function is used with lower.limits=0, alpha=0, lambda=0, intercept=FALSE. These settings perform a linear regression where all coefficients are non-negative, and with no regularization and no intercept term.

## Reference Data

### BLUEPRINT Reference Dataset

35 gene counts files were downloaded from the BLUEPRINT database, all collected from venous blood [18]. This included entries for CD14-positive, CD16-negative classical monocytes (5 samples), CD38-negative naive B cells (1), CD4-positive, alpha-beta T cell (8), central memory CD4-positive, alpha-beta T cell (2), cytotoxic CD56-dim natural killer cell (2), macrophage (4), mature neutrophil (10), and memory B Cell (1). When two or more transcripts appeared for a single gene, the transcript with the highest average expression was selected and others were excluded. Genes with no detected expression in any sample were also excluded, and then each sample was quantile normalized. Samples generally clustered by cell type, but we excluded one CD4-positive alpha-beta T cell. Replicates for each cell type were then collapsed into a single entry by taking the median value for each gene.

### ENCODE Reference Dataset

106 transcript quantification files were downloaded from the ENCODE database [19]. These included all RNA-seq experiments collected from adult primary cells, excluding four with warnings. Warnings indicated that three samples suffered from low replicate concordance and one sample from low read depth, and these samples were excluded. All samples were processed by the Gingeras Lab at Cold Spring Harbor and mapped to GRCH38.

The samples were quantile normalized and clustered. In cases where multiple transcripts were measured for a single gene, the expression of that gene was calculated as the sum of all transcripts. At this time, 18 additional samples were excluded as they did not cluster with their replicates. Based on sample descriptions and data clustering, we found that the remaining 88 samples represented 28 unique cell types. We produced an expression profile for each cell type by merging all samples of that cell type via median average. For example, a cluster of 19 samples were labelled as endothelial cells (collected from various body locations) and were merged into a single entry termed canonical endothelial cells. This dataset spans a wide range of stromal cell types (e.g. smooth muscle, fibroblast, epithelial), but contains only a single entry for blood cells, which are labelled mononuclear cells.

We also combined the ENCODE and BLUEPRINT reference matrices into a single reference matrix, which we call BlueCode. We combined, then quantile normalized, the columns of both matrices. Possible batch effects in this combined matrix have not been fully evaluated.

### 10x Reference Dataset

We obtained single cell expression data for nine varieties of immune cells from the 10x website [20]. This included at least 2446 cells for each cell type, and at least 7566 cells for all cells other than CD14 monocytes. For each cell type, expression values for all cells were mean averaged to form an expression profile.

### Tabula Muris Reference Dataset

We downloaded from the Tabula Muris single cell data for 12 clusters of mouse cell types. For each cluster, we averaged all cells of that cluster to produce a reference profile for the corresponding cell type.

### Other Reference Datasets

Other datasets used in this project were obtained from their corresponding publications or GEO repositories. This includes a reference matrix of human skin signatures, the Human Body Atlas, the Human Primary Cell Atlas, LM22, ImmunoStates, the Mouse Body Atlas, and ImmGen [14–17,21,23,24].

### Skin Diseases Data

We obtained expression data from 21 skin biopsies, collected from human patients with a variety of skin diseases. These data originally came from a wide range of sources and platforms, and were compiled into a single dataset by previous work [35].

### GTEx Data

GTExX data for 17,382 samples were obtained from the GTExX database (https://gtexportal.org/). We ran GEDIT on all samples three times, each time using a different reference matrix (BlueCode, the Human Primary Cell Atlas, and Skin Signatures). For each cell type, we calculated our initial estimate as the median estimate across the three sets of predictions (or fewer, if that cell type is missing from one to two of the reference matrices). Lastly, for each sample we divided the vector of predictions by its sum, such that the final predictions sum to 100%.

## Multi-Tool Performance Evaluation

### *In Vitro* Immune Cell Mixture

Combinations of six immune cells (Neutrophils, Monocytes, Natural Killer Cells, B cells, and CD4 and CD8 T Cells) were mixed together and sequenced using an affymetrix array. Whole blood from healthy human donors was supplied with informed consent through a sample sharing agreement with the UCLA/CFAR Virology Core Lab (grant number 5P30 AI028697). CD4+ T cells, CD8+ T cells, B cells, and NK cells were isolated using Stem Cell Technologies (Vancouver, BC, Canada) RosetteSep negative selection. Neutrophils were positively selected through the EasySep approach, according to the manufacturer’s specifications. Cells were then counted by hemocytometer and added at defined percentages to a total cell count of two million cells to create six different mixtures. Subsequently cells were processed for RNA isolation by AllPrep DNA/RNA. Illumina HT12 BeadChip microarray was performed by the UCLA Neuroscience Genomics Core. Data was normalized by quantile normalization through R ‘normalize.quantiles’ function (R Core Team, 2013).

### RNA-seq Mixtures Used for Tool Evaluation

We also obtained two datasets used in a recent benchmarking study [30]. The first dataset is composed of three RNA-seq samples, each with two technical replicates that represent biopsies of ovarian cancer ascites [32]. The second dataset is composed of RNA-seq collected from the blood of healthy individuals, some of whom recently received an influenza vaccine [31]. These data were downloaded from the GitHub for the benchmarking paper, which also contained FACS estimates for six cell types for the ascites data (B cells, dendritic cells, NK cells, T cells, macrophages, neutrophils) and five cell types for the blood data (B cells, dendritic cells, T cells, monocytes, natural killer cells). However, since dendritic cells were never present at more than 3.5% abundance, we did not evaluate performance for this cell type.

### Tools

We installed and ran GEDIT, CIBERSORT, DeconRNASeq and dtangle on the hoffman2 computational cluster at UCLA. xCell was run using the online interface at https://xcell.ucsf.edu/. The default choice for genes signatures (xCell =64) was used. The RNA-seq option was selected for the 2 RNA-seq datasets (blood and ascites), but not for the *in vitro* dataset, which was sequenced on microarray.

xCell produces 67 output scores, seven of which were used in this study. These were the entries labelled “B-Cells”, “Macrophages”, “Monocytes”, “NK cells”, “Neutrophils”, “CD4+ T cells” and “CD8+ T Cells”. As suggested by the xCell authors, the outputs for CD4 and CD8 T cell subtypes were summed to produce a final output for total T cells.

### Reference Data

We evaluated the performance of the four reference-based tools (GEDIT, CIBERSORT, DeconRNASeq and dtangle) using each of four choices of reference matrix (LM22, ImmunoStates, BLUEPRINT, and the Human Primary Cell Atlas).The BLUEPRINT and Human Primary Cell Atlas reference matrices differ from ImmunoStates and LM22 in that they contain tens of thousands of genes, many of which should not be considered signature genes. This contrasts to ImmunoStates and LM22; each reference matrix contains fewer than 600 genes, which have been specifically identified as signature genes by previous work [14,21]. We include both forms of reference matrices in order to evaluate the input requirements of the tools studied.

Depending on the choice of reference matrix, reference-based tools often produce multiple outputs for some cell types, each representing a cell sub-type. This includes B cells (naïve and memory), Monocytes (CD14 and CD16), NK cells (resting and active) and T cells (many subtypes including varieties of CD4 and CD8). In each case, the outputs for each sub-type were summed in order to produce a total score for each greater cell type.

## Discussion

GEDIT is an expression-based cell type quantification tool that offers unprecedented flexibility and accuracy in a wide variety of contexts. Using both simulated and experimental data, we demonstrate that GEDIT produces high-quality predictions for multiple platforms, species, and a diverse range of cell types, outperforming other tools in many cases. We include in the software package a comprehensive library of reference data, which facilitates application of GEDIT to a wide range of tissue types in both human and mouse. GEDIT can also accept reference data supplied by the user, which can be derived from bulk RNA-seq, scRNA-seq, or microarray experiments. GEDIT represents a competitive addition to the suite of existing tissue decomposition tools while maintaining flexibility and performance robustness.

As part of this project, we perform a study in which we compare the performance of several deconvolution tools using multiple metrics. Unlike previous evaluation studies, we explore the effect of reference choice by running tools multiple times with reference data from different sources. Choice of optimal reference has a large impact on the accuracy of many tools, but GEDIT provides robust performance and accurate estimates for many possible reference choices. While all efforts were taken to perform this comparison in an unbiased manner, the authors note that development of the tool was still underway when the first comparisons were made. All code and inputs used to reproduce this study are included in the github (https://github.com/BNadel/GEDIT), with the exception of CIBERSORT code, which is limited by copyright.

The high performance of GEDIT is due to two key innovations. Firstly, signature gene selection by information entropy serves to select genes that are the most informative for deconvolution. Secondly, the row scaling step, which aims to equally weight all signature genes into the final estimate, even those with comparatively low expression. In addition, the flexibility of GEDIT and the diverse set of reference matrices we provide allows GEDIT to be easily applied in a wide range of circumstances.

The output of GEDIT represents the fraction of mRNA originating from each cell type. This is an effective measure of the transcriptional contribution of each cell type in a mixture. However, in cases where some cell types consistently produce more or less mRNA per cell, this measure may not represent cell counts. Data capturing the average mRNA content per cell is becoming more widely available in the form of single cell experiments and could in principle be used to convert our fractions into cell counts.

When extensively applied to several large public datasets, GEDIT produces predicted cell type fractions that conform with biological expectations. When used to decompose skin biopsies, keratinocytes are found to be the most abundant cell type. Variations in the abundance of other cell types conform to expected immune responses across diseases. Similarly, cell type predictions of GTEx samples are concordant with our expectations of the dominant cell types across tissues. Schwann cells, keratinocytes, adipose cells, and immune cells are found to be most abundant in nerve, skin, adipose tissue, and blood, respectively.

Single cell RNA-seq is an emerging approach to study the composition of cell types within a sample. Due to biases associated with the capture of different cell types, these methods are not always capable of accurately quantifying cell type populations [8]. However, the pure reference profiles produced by existing methods can be used by GEDIT to generate accurate estimates of cell type populations. Thus, GEDIT circumvents some of the biases associated with the preparation of samples for both scRNA-seq and FACS. GEDIT is freely available, and therefore an extremely economical option for researchers, particularly those who profile expression data for other purposes.

GEDIT produces accurate results when tested on mixtures of human immune cells. Compared to other tools, GEDIT produces the lowest error in majority of scenarios in the studied mixtures. GEDIT provides increased flexibility over previously developed tools, as we provide a set of reference matrices for varied cell types for both mouse and human datasets.

GEDIT provides unique advantages to researchers, especially in terms of cell type, species and platform flexibility, and constitutes a useful addition to the existing set of tools for tissue decomposition. Our efficient decomposition methodology has been extensively optimized and we find that it performs robustly across a broad range of tissues in both mouse and human datasets. Our future work will extend reference matrices to facilitate application of GEDIT on varied bulk gene expression datasets.

## Availability of Source Code and Requirements

⍰ Project name: GEDIT
⍰ Project Home Page: https://github.com/BNadel/GEDIT
⍰ Programming Languages: Python 2.0, R
⍰ Other requirements: numpy, glmnet
⍰ Operating Systems: Linux
⍰ License: MIT

## Availability of Data and Materials

All data used in this paper are freely available on GitHub (https://github.com/purebrawn/GEDIT), as well as their original sources. Code for DeconRNASeq was obtained as an R package from the CRAN repository. Code for CIBERSORT was obtained by requesting it via the web portal (https://cibersort.stanford.edu/download.php), and code for dtangle from the project’s GitHub page (https://github.com/gjhunt/dtangle).

Reference data is also available from their original sources. Most datasets can be found on project website pages or from public databases. These include BLUEPRINT (http://www.blueprint-epigenome.eu/), ENCODE (https://www.encodeproject.org), the Human Primary Cell Atlas (http://biogps.org/dataset/BDS_00013/primary-cell-atlas/), LM22 (http://cibersort.stanford.edu/%20or%20GEO:GSE65136), 10x Genomics (https://support.10xgenomics.com/single-cell-gene-expression/datasets), Tabula Muris (https://tabula-muris.ds.czbiohub.org/), the Mouse Body Atlas (GEO:GSE10246), and ImmGen (http://www.immgen.org/Databrowser19/DatabrowserPage.html). Some reference matrices were obtained as supplementary files from the publications listed in Table 1.

Expression values for the blood and ascites RNA-seq datasets were obtained from the GitHub repository https://github.com/grst/immune_deconvolution_benchmark, and are also available at at https://figshare.com/s/711d3fb2bd3288c8483a and GEO: GSE64655). The *in vitro* mixture of immune cells was prepared by our lab, and available on our GitHub page.

## Acknowledgments

We acknowledge the Biomedical Big Data Grant (5T32LM012424-03) for supporting Brian Nadel during the course of this research. We also acknowledge the Bruins-in-Genomics Summer Undergraduate Research Program for supporting Hannah Waddel and Misha Khan during the summer of 2017, when they contributed to this work. We also thank Lana Martin for her help with editing and proofreading the manuscript.

