## Supplementary Materials for "The Gene Expression Deconvolution Interactive Tool (GEDIT): Accurate Cell Type Quantification from Gene Expression Data"

**Synthetic Mixture Generation**

The deconvolution of synthetic mixtures using only a single matrix (to both generate the mixtures, and serve as a reference) is a trivial problem. In this context, the linear regression will always return the exact (or nearly exact) input proportions. Moreover, this is a poor simulation of real world data, as in reality the expression profile of any given cell type will vary to some extent between experiments. The mixtures submitted by the user will often be from different platforms than the reference data, and, in particular, cross-platform effects cannot be simulated using a single matrix. Therefore, in order to more meaningfully evaluate the performance of deconvolution, we used a separate matrix to produce mixtures from the one used as a reference.

Using distinct reference and mixture-generating matrices requires that we match cell types between the two matrices. Matching cell types across references is a non-trivial problem, as equivalent cell types may be labelled differently, and identically labelled cell types may not be equivalent. To address this problem, we defined the following procedure for identifying pairs of equivalent cell types between two reference matrices:

1. Joint quantile normalize the matrices, then log transform them
2. Calculate the Pearson correlations between each cell in the first matrix and each cell in the second matrix
3. Pair cell types that are more highly correlated with each other than with any other cell type in the reference
4. Manually exclude cell pairings with mismatching descriptions

Using this procedure, we identified 5 pairings of reference matrices that can be used for the generation of synthetic mixtures (Table 2). Since simulations can be done in both directions for each pair, this represents 10 possible choices of a mixture generating matrix and a reference matrix.

| Matrix1 | Matrix2 | Number of Cell Types | Platforms |
| --- | --- | --- | --- |
| BluePrint | Human Primary Cell Atlas | 5 | RNASeq to Affymetrix U133 Microarray |
| BluePrint | 10x Single Cell | 4 | Bulk RNASeq to SC RNASeq |
| BluePrint | Skin Signatures | 6 | RNASeq to Affymetrix/Illumina HT-12 Microarray |
| Human Primary Cell Atlas | Skin Signatures | 10 | Affymetrix U133 Microarray to Affymetrix/Illumina HT-12 Microarray |
| 10x Single Cell | Skin Signatures | 4 | SC RNASeq to Affymetrix/Illumina HT-12 Microarray |

Supplementary Table 1. Pairs of reference matrices used to generate synthetic mixtures.

For each of these 10 pairs of matrices, 1,000 cell type proportions were generated randomly. Specifically, a cell type was selected at random and assigned a weight between 0 and 1.0 (randomly sampled from the uniform distribution). Next, one of the remaining cell types is randomly selected and assigned a weight between 0.0 and the remaining weight (1.0 minus the sum of weights already assigned). This is repeated until the final cell type, which is assigned all the remaining weight.

The final simulated expression profile is produced by summing the expression profiles of each cell type, multiplied by the simulated weight. We believe this procedure produces biologically reasonable mixtures, as they are composed primarily of a small number of cell types, with many other cell types present at low levels.

**Signature Gene Selection**

We have tested a total of 6 signature gene scoring algorithms; Entropy, fsDiff, fsRatio, meanDiff, meanRatio, and Zscore. For a given gene, these algorithms take as input the vector of expression values across all cell types, and return a score. Each gene is a candidate signature gene for the cell type in which it is most highly expressed, and only genes with the highest signature scores are accepted. The number of genes selected is determined by the NumSigs parameter, which is by default set to 50.

One scoring approach is to compare the highest observed expression value to the mean of all other expression values. This comparison can be performed by division or subtraction (MeanDiff and MeanRat). Alternately, these same comparisons can be made between the highest observed expression value, and the second highest observed value (fsDiff and fsRat). The Zscore method is calculated the same way as MeanDiff, except that it is divided by the standard deviation of the expression vector.

When run on 10,000 simulated mixtures, selecting genes by entropy produced the lowest maximum, mean, and upper quartile error (Figure 2A). We therefore use entropy as the default setting, but allow the user to select any of the other 5 scoring methods. Using entropy has the potential to select genes that are highly expressed in 2 or more cell types, and lowly expressed in the rest. While these genes are not unique to a single cell type, they can still offer valuable information for deconvolution.

**Number of Signature Genes (NumSigs, MinSigs)**

GEDIT’s second parameter is the number of signature genes that are selected per cell type. On simulated data, any number of signature genes between 40 and 200 produce near-optimal results (Figure 2B).

We provide an option that allows more signature genes for some cell types than others. In this scheme, both an average and a minimum number of signature genes are specified by the user (NumSigs and MinSigs, respectively). For each of N cell types present in the reference, MinSigs genes are selected that are maximally expressed in that cell type. However, a total of N*NumSigs genes are selected, and the remaining N*(NumSigs-MinSigs) genes are simply those with the highest score, regardless of the cell type in which they are maximally expressed.

On simulated data, we found that adjusting the MinSigs parameter had minimal effect on predictions (Figure 2D), and by default GEDIT sets MinSigs equal to NumSigs.

**Row Scaling**

The extent of row scaling is controlled by the row scaling parameter, with allowed values between 0.0 and 1.0. At 1.0 a gene with 10x higher expression will have 10x the influence (as if no row scaling were performed). At a value of 0.0, all genes have equal influence. In simulated experiments, a row scaling value of 0.0 produced the lowest mean error, substantially improving accuracy (Figure 2C). Values outside the natural range of 0.0 to 1.0 produce high error, as well (data not shown).

**Multi-Tool Comparison**

Here, we include figures presenting the errors and correlations for each mixture used (and each cell type in those mixtures). This contrasts Figure 4, where values are considered for either each cell type (including all datasets), or each dataset (including all cell types).

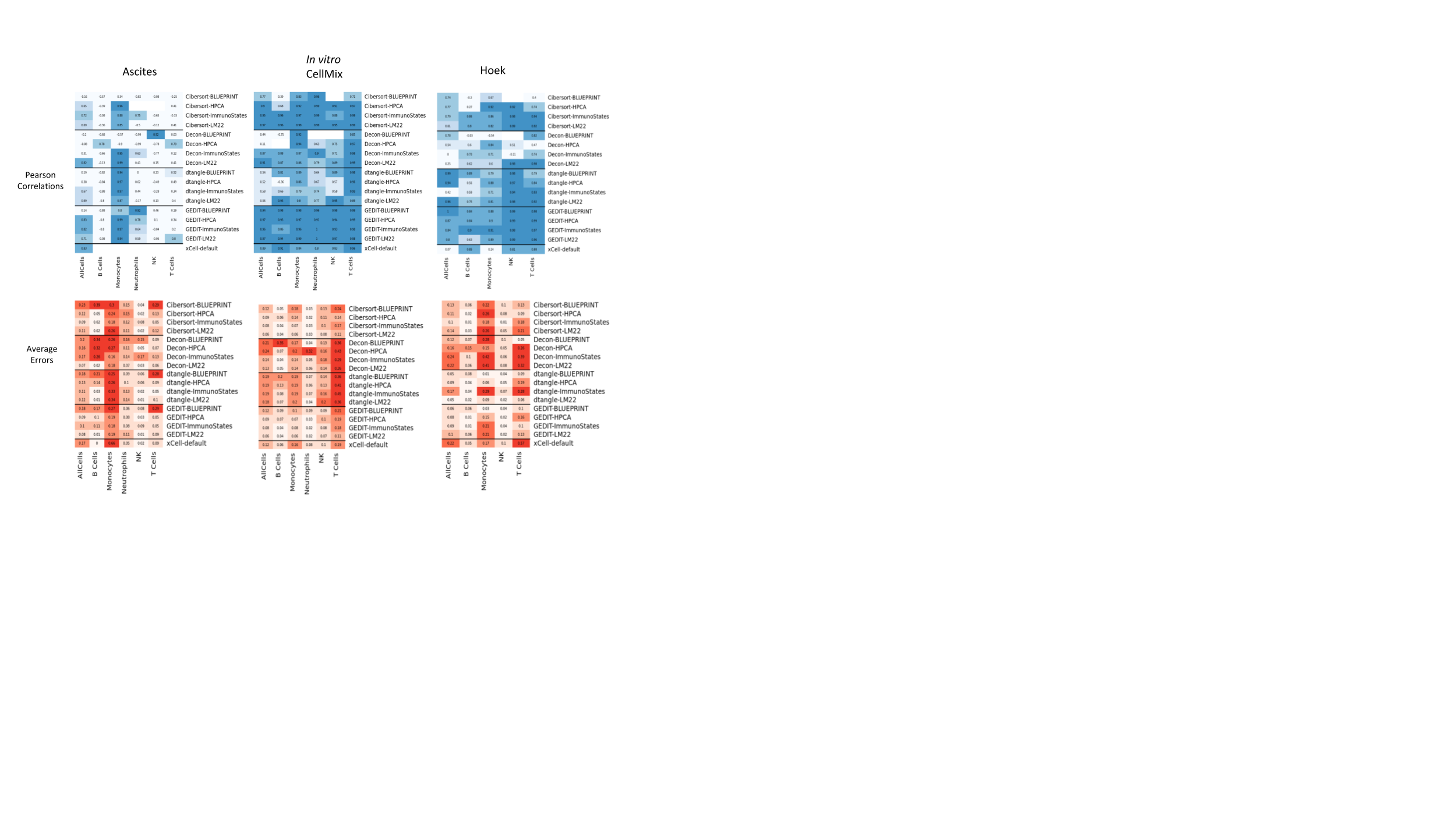
Supplementary Figure 2. Error and correlation values when benchmarking is performed on 3 datasets (ascites, Hoek, and *in vitro* cell mixtures) using each of 5 tools (CIBERSORT, DeconRNASeq, dtangle, GEDIT, and xCell) and each of 4 possible reference matrices (BLUEPRINT, HPCA, ImmunoStates, LM22)

We also perform additional comparisons between bulk deconvolution tools using simulated data. Firstly, we apply the tools to a set of simulated mixtures created using single cell data from the human pancreas. Specifically, we use the framework implemented as part of the SCDC project [[[1]](https://paperpile.com/c/noZFqs/yLaW)] to produce 100 simulated pseudo-bulk mixtures from a single cell experiment on human pancreas [[[2]](https://paperpile.com/c/noZFqs/Esh9)]. We utilize data from a separate single cell human pancreas experiment to produce reference profiles [[[3]](https://paperpile.com/c/noZFqs/tfUe)]. This is done by calculating the average expression profile of all cells of each cell types (alpha, beta, gamma, delta).

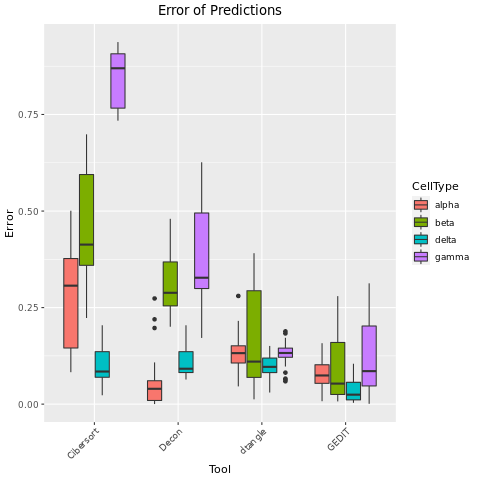

Supplementary Figure 3. Distribution of error values between predicted and actual fractions when deconvolution is applied to a set of 100 simulated pancreatic samples. Simulated samples are created using data single cell data from human pancreas [[[2]](https://paperpile.com/c/noZFqs/Esh9)], and separate single cell data is used to create a bulk reference matrix [[[3]](https://paperpile.com/c/noZFqs/tfUe)].

Relative to the other 3 tools, GEDIT performs well on this dataset. It produces the lowest average error across all cells, and also the lowest average error for beta, delta, and gamma cells. DeconRNASeq produces the lowest error for alpha cell predictions, but performs poorly when predicting beta and gamma cells.

**Runtime Analysis**

We also perform an experiment in which we vary the number of inputs submitted to each tool, and record the computational time. We randomly select 100, 200, 500, 1000, and 2000 samples from the GTEx database, and run each locally available tool on the hoffman2 cluster at UCLA. We repeat this experiment six times, such that six datapoints are available for each batch size. Each tool was provided 16 gigabytes of RAM, and CPU time was recorded. CIBERSORT is the slowest to run at every input size. The other 3 tools consistently produce results within 25 minutes, even for the largest number of inputs (2000).
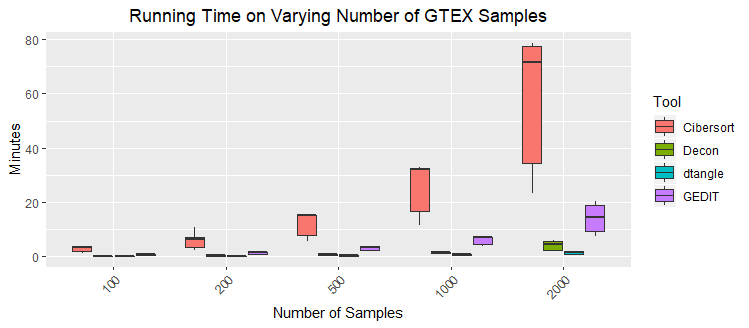

Supplementary Figure 3. CPU time for deconvolution tasks to complete when applied to inputs of varying size. Inputs are created by selecting varying numbers of samples from the GTEx database. For each input size, samples were randomly selected six times and deconvolution performed a total of 24 times, once for each of 4 tools. The LM22 matrix is used as a reference profile.

**Deconvolution of Human Skin Diseases**

We show here the predictions of each tool when applied to a series of 21 samples from various human skin diseases. The reference used here is the Skin Signatures matrix.

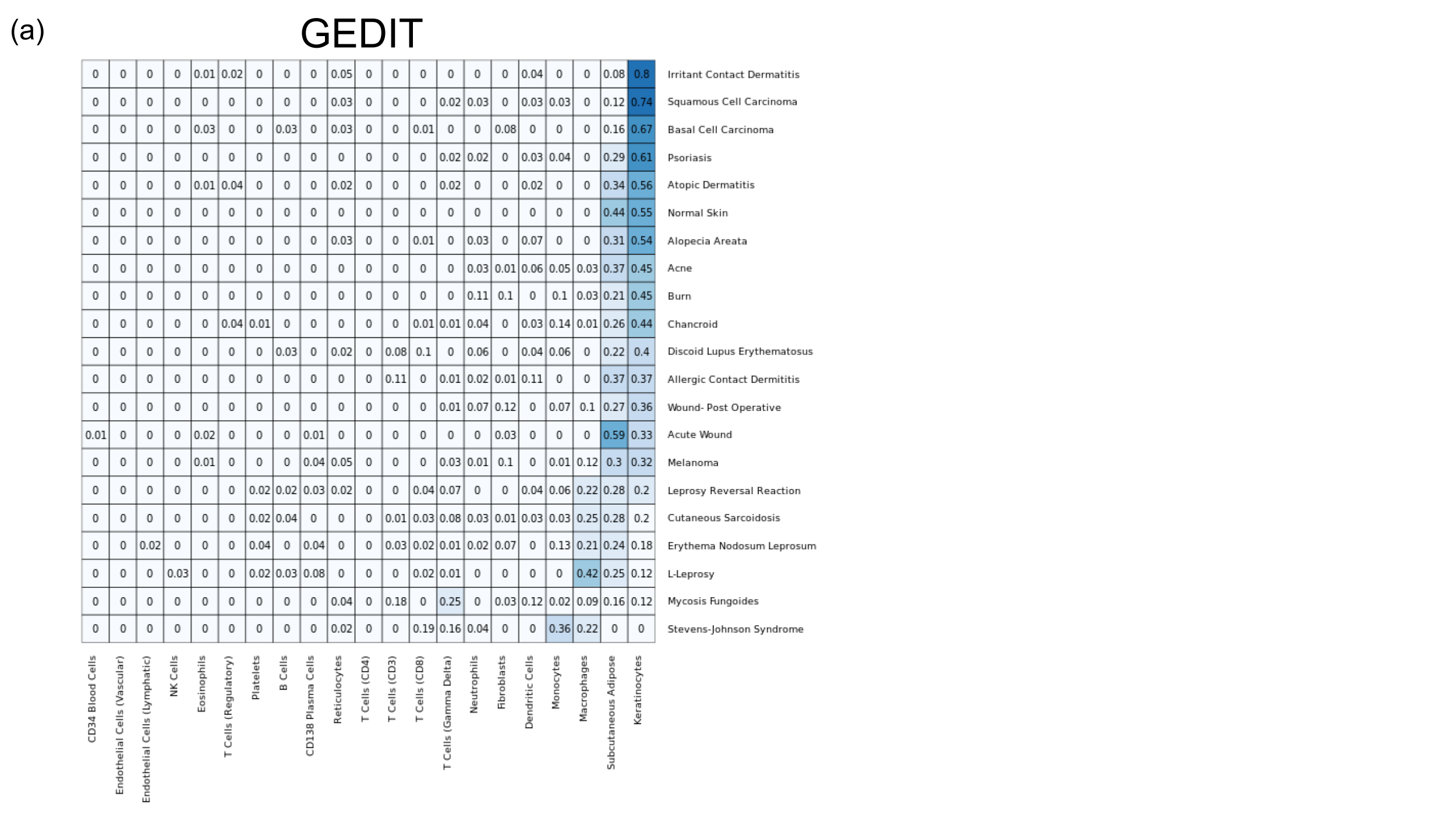

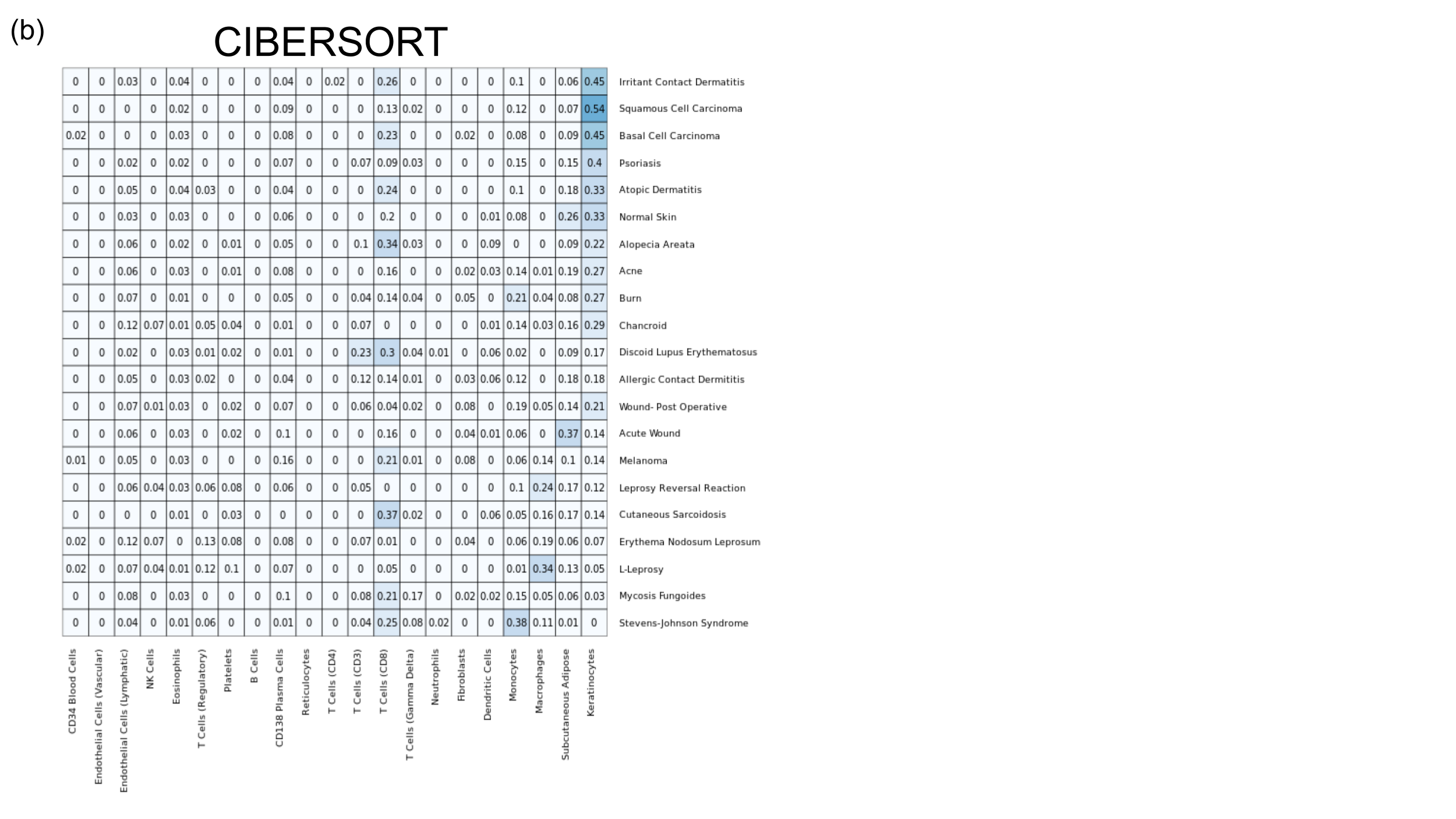

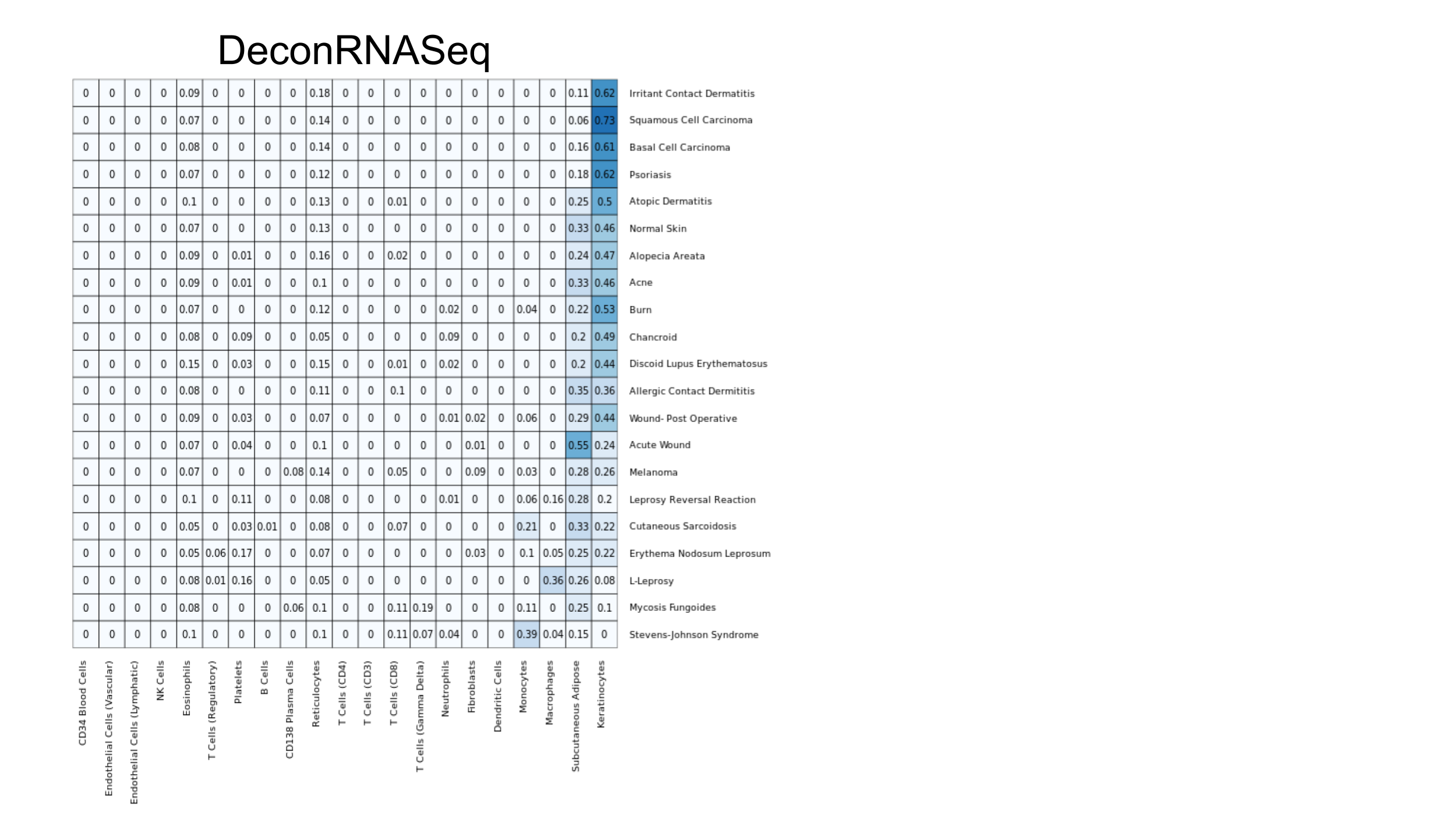

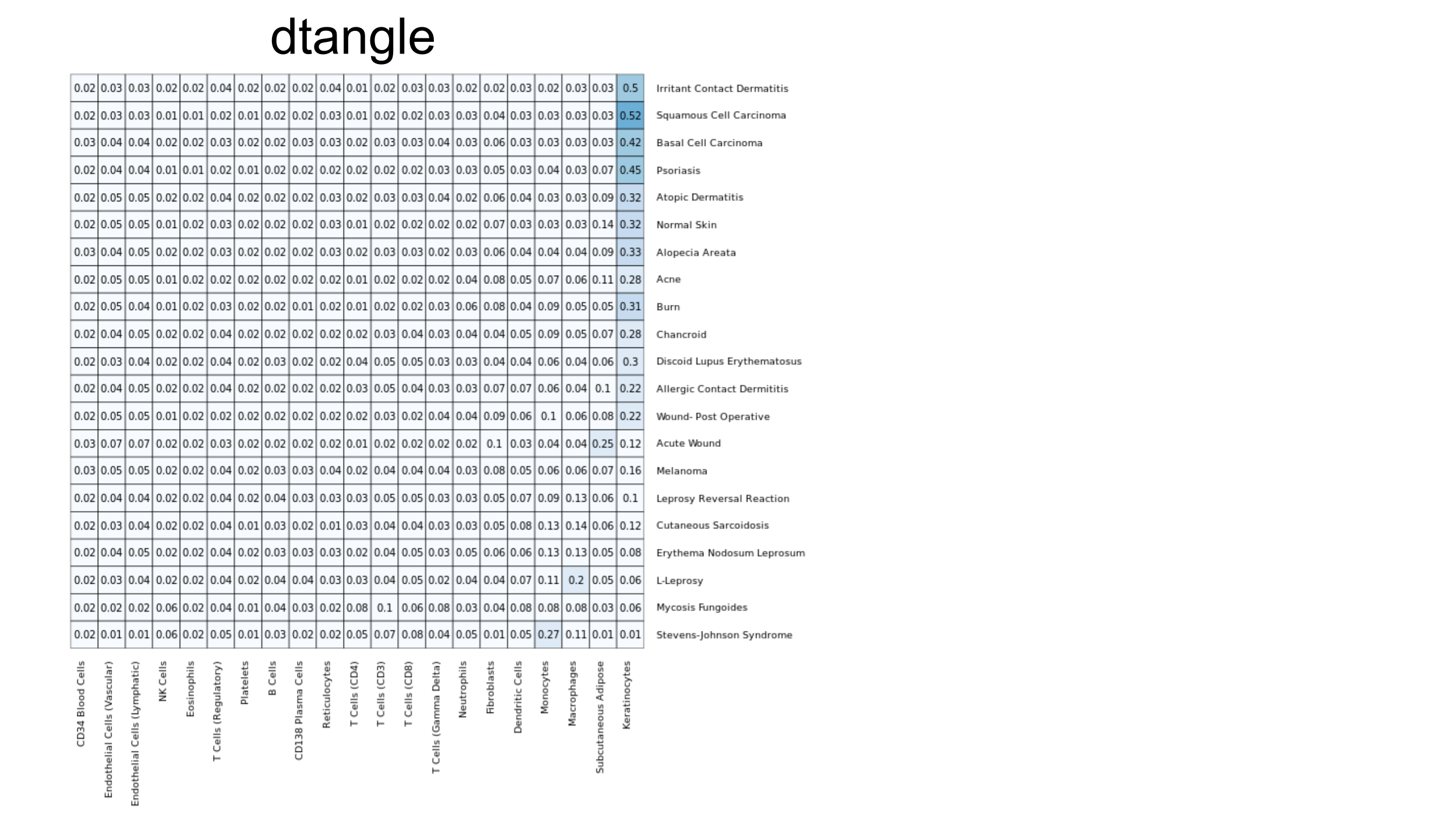

Supplementary Figure 4a-d. Predicted cell type fractions for 21 skin samples using each of 4 tools. Samples represent microarray expression data from 21 biopsies of skin diseases, and the Skin Signatures matrix is used as a reference [[[4,5]](https://paperpile.com/c/noZFqs/KSlg+c8NZ)].

**Deconvolution of the Mouse Body Atlas**

We have also produced predictions of cell type content for 30 samples from various body sites in mouse, as obtained and published in the Mouse Body Atlas ([[6]](https://paperpile.com/c/noZFqs/9vDd)**)**. The reference matrix used here Tabula Muris is constructed from the Tabula Muris ([[7]](https://paperpile.com/c/noZFqs/FKXV)).

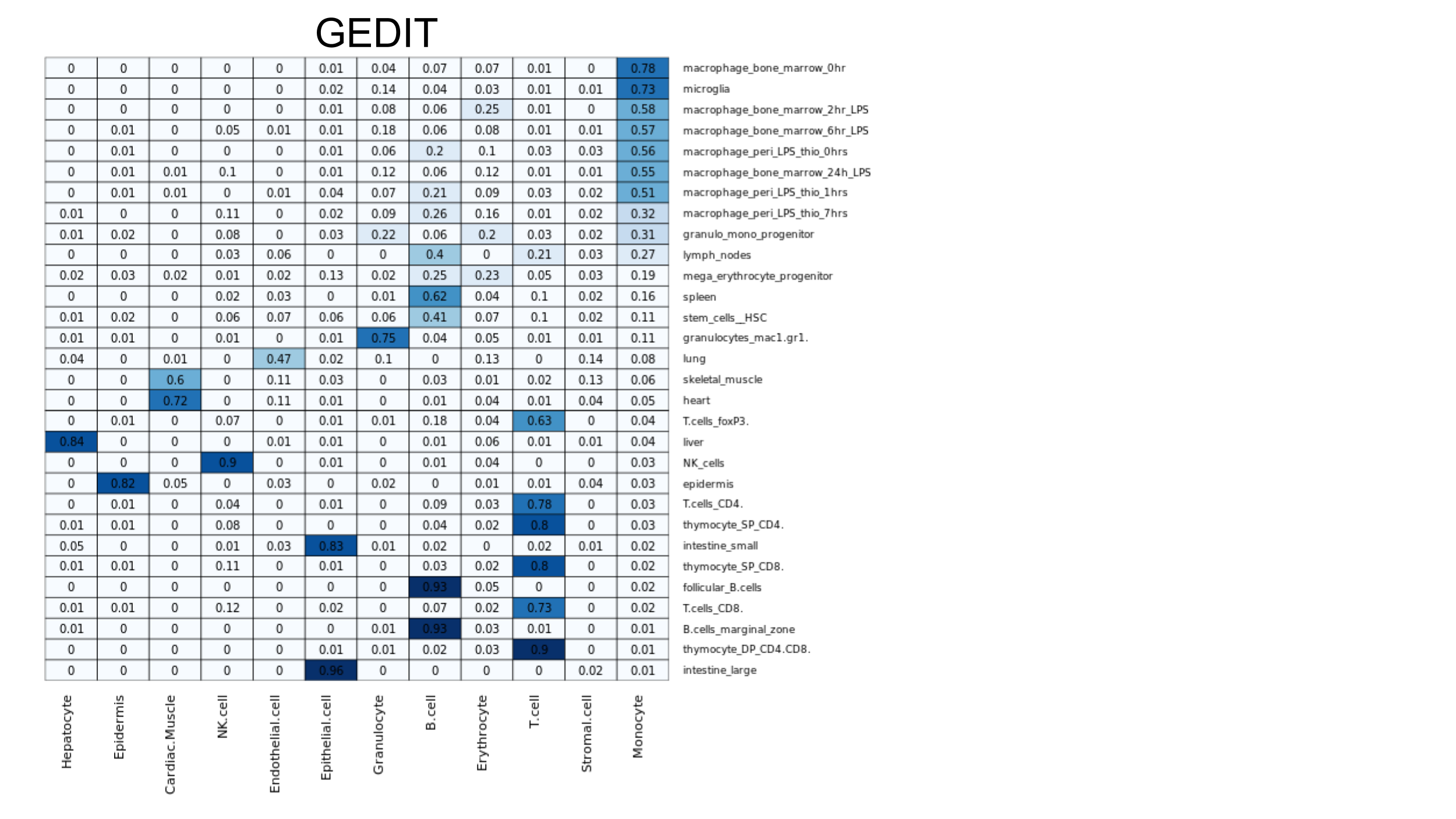

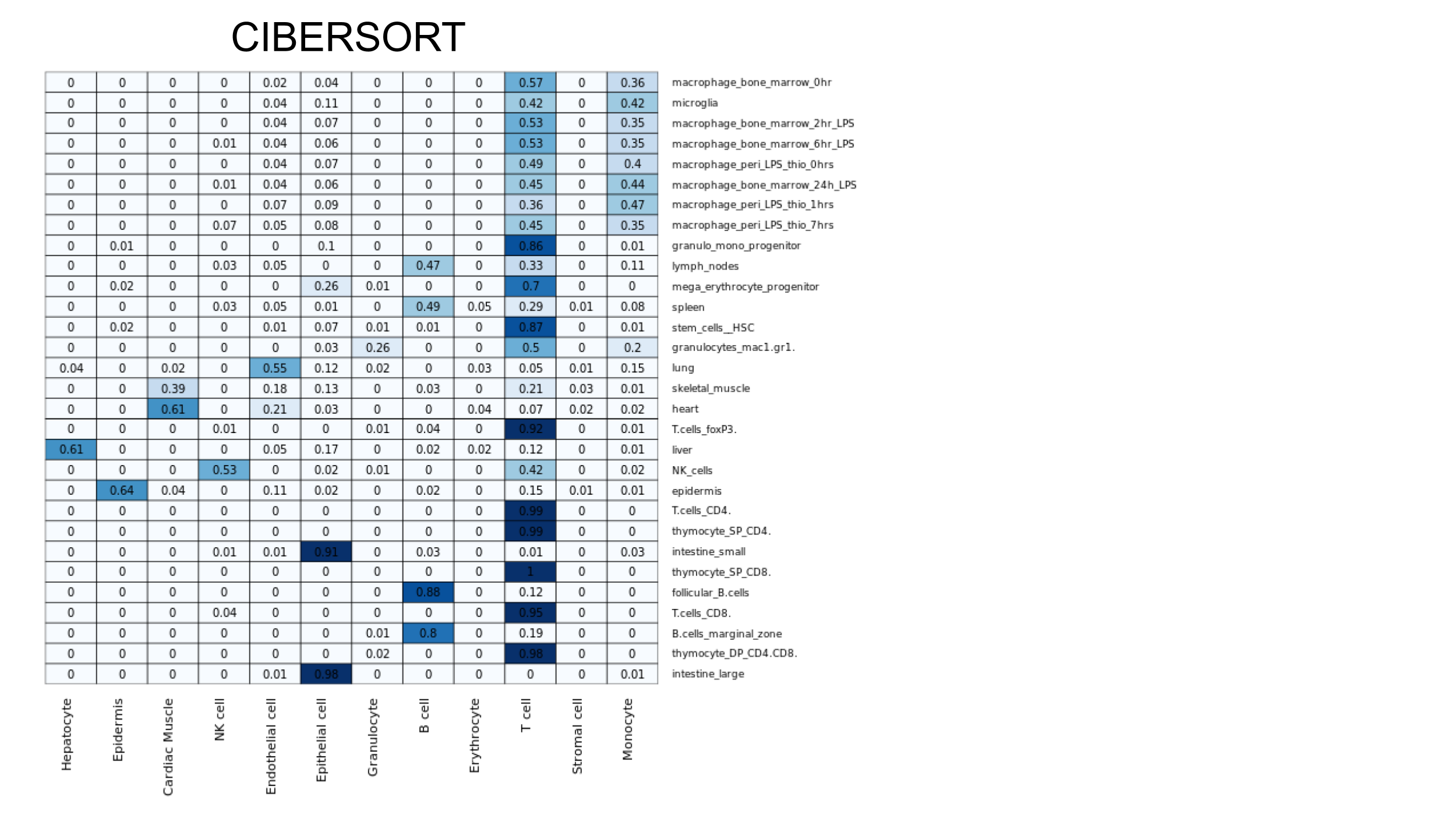

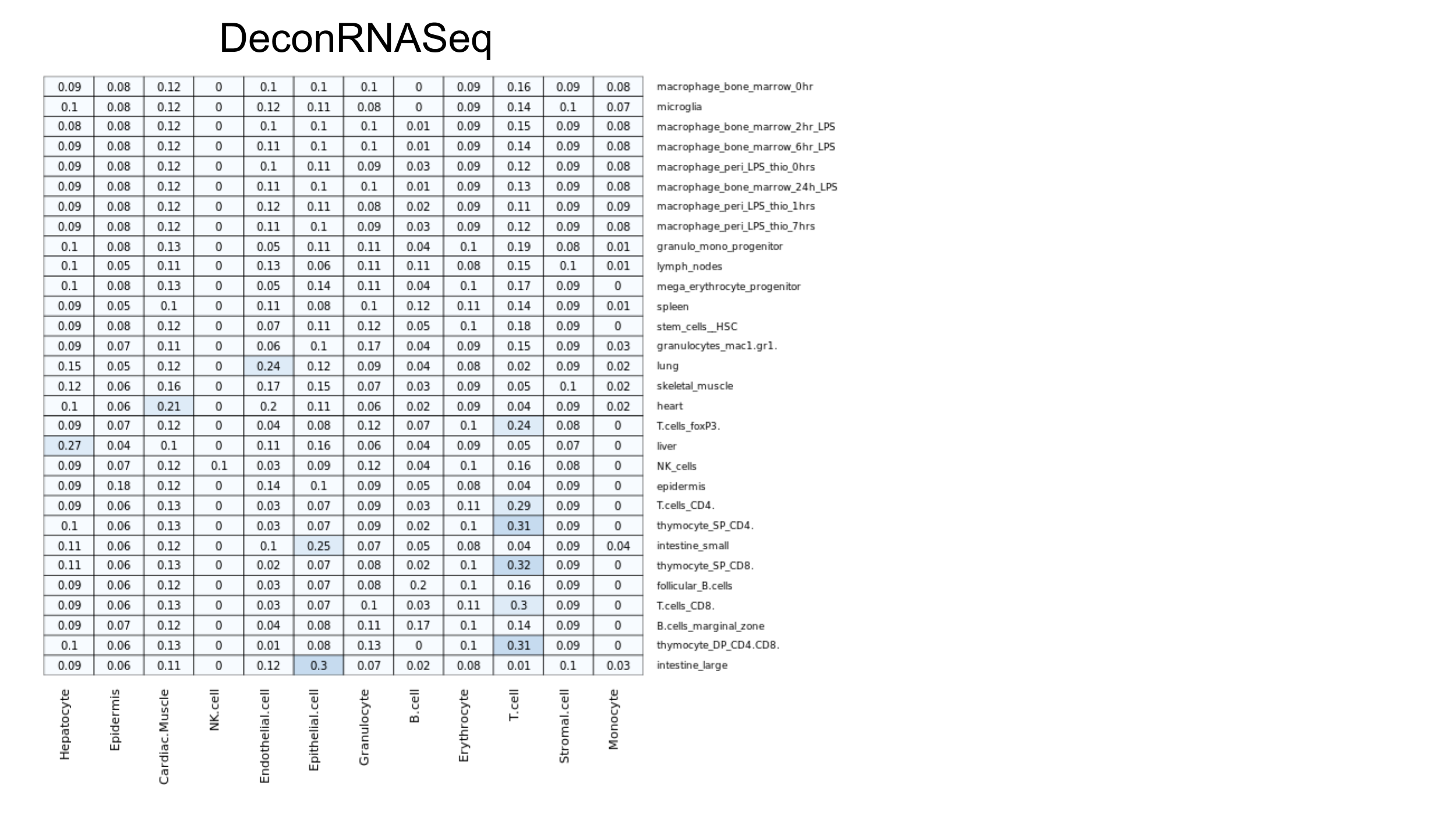

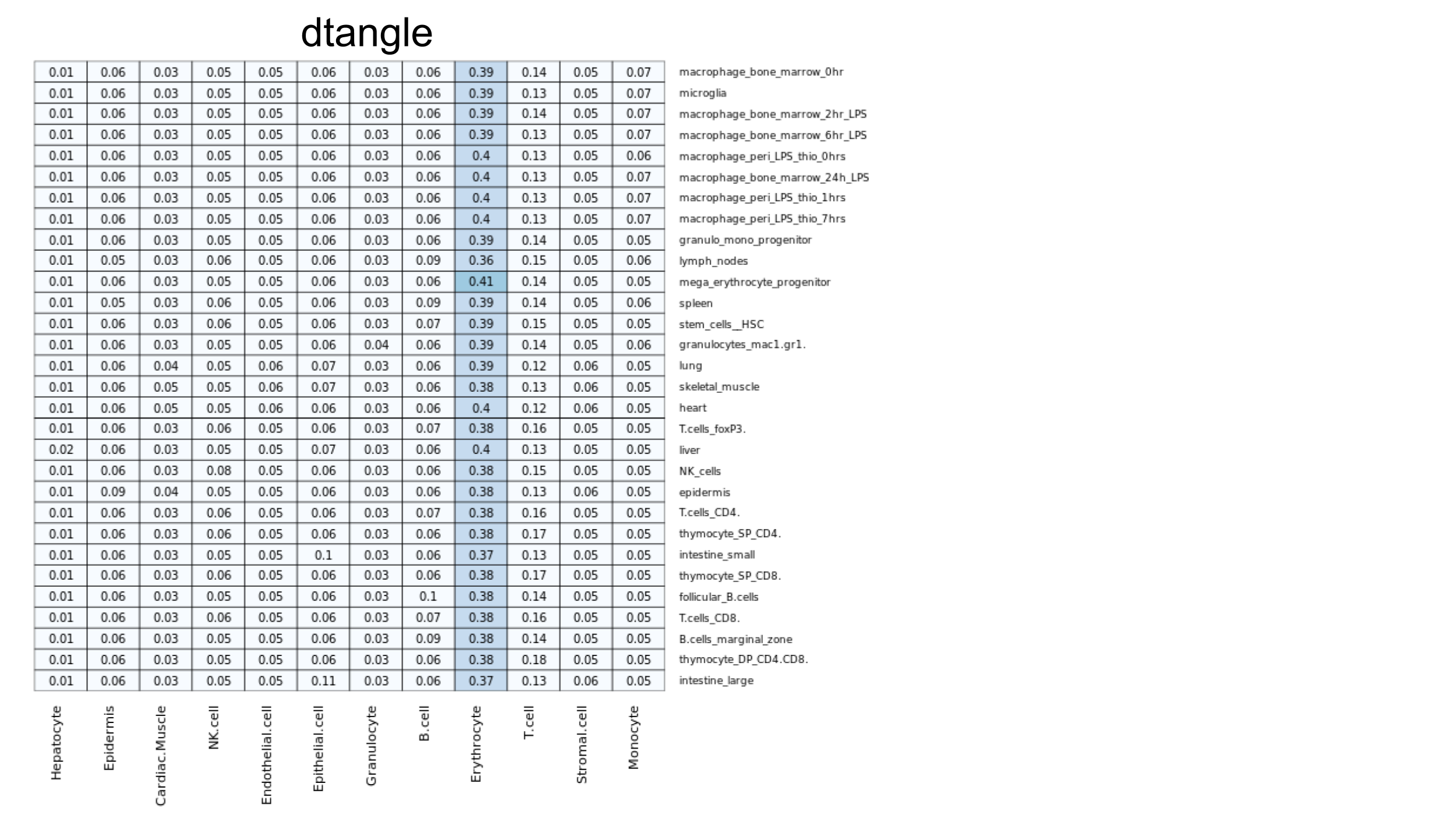

Supplementary Figure 5a-d) Predicted cell type composition of 30 samples from the Mouse Body Atlas[[6]](https://paperpile.com/c/noZFqs/9vDd). Samples were profiled using an Affymetrix U133A/GNF1H microarray. Single cell data from the Tabula Muris was averaged for each cell type to create a bulk reference [[7]](https://paperpile.com/c/noZFqs/FKXV).

**Deconvolution of GTEX Database**

We used GEDIT to estimate the cell type proportions of 10 cell types for 17,382 samples in the GTEX database. Since no single reference matrix contained all 10 cell type, we took an approach utilizing several reference matrices (Supplementary Figure 1). First, we combined the BLUEPRINT reference matrix (which contained only immune cells) with the ENCODE reference matrix (containing mostly stromal cells). We did this by concatenating the columns then quantile normalizing, and refer to the resulting matrix as “BlueCodeV1.0”. We then ran GEcd DIT deconvolution on the entire GTEX database three times; once using BlueCodeV1.0 as the reference, once using the Human Primary Cell Atlas, and once using the Skin Signatures matrix. For each predicted fraction in each sample, we took the median value of the 1-3 predictions produced. Lastly, we divided the predictions of each sample by their sum, such that predictions summed to 1.0. These values were used as final estimates for the fraction of each cell type in each sample.

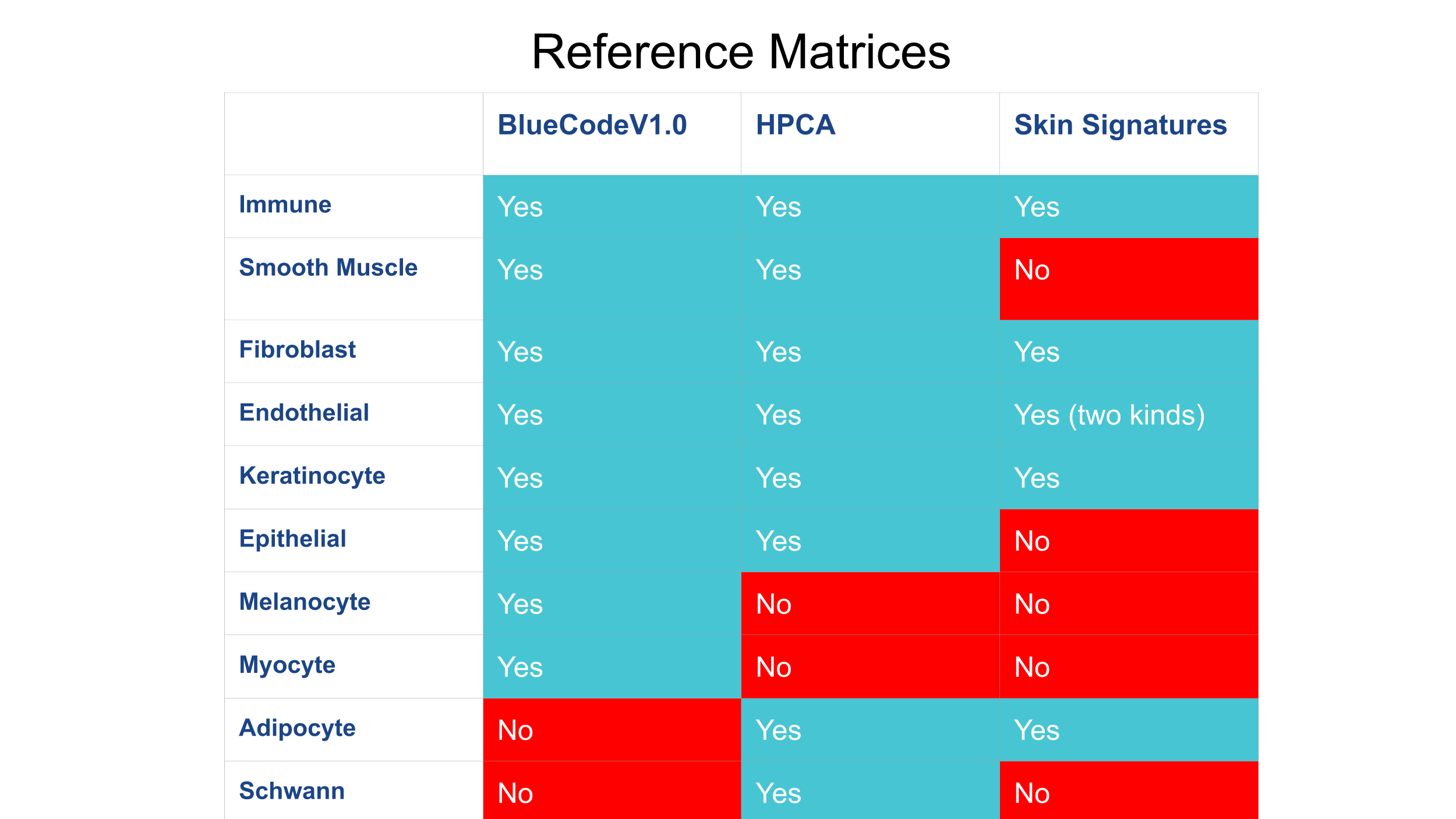

Supplementary Figure 6. Cell types present in the 3 reference matrices used to predict cell type fractions of GTEX samples

We also explore the effects of merging these datasets into a single reference matrix prior to performing deconvolution. We combine 27 fields from BluePrint and ENCODE with 8 fields from the Human Primary Cell Atlas. The 8 HPCA cell types were absent in the other 2 references, and included Adipocytes. Schwann Cells, CD34+ Cells, Platelets, Dendritic cells (BDCA1+) and gamma delta T cells. Keratinocytes were also included, as the keratinocytes available from ENCODE are specifically “hair follicular keratinocytes” and may have a different expression profile compared to keratinocytes from other locations.

Clustering reveals that batch and/or platform effects are present in this combined matrix. Cell types from HPCA do not cluster with similar cell types from the BLUEPRINT and ENCODE databases. When using this reference to deconvolute the GTEx database, some predictions correspond to known biology (e.g. keratinocytes in skin, B Cells in spleen), but others do not (e.g. keratinocytes in testes and brain, myometrial cells in liver). In this case, it appears that combining reference data from separate platforms is problematic for deconvolution. Instead, we use the approach described above where we run deconvolution using each reference matrix independently and merge the predictions afterward. Both approaches have limitations, and as more data becomes available in the future, creating a comprehensive reference from single cell data will likely produce superior results.
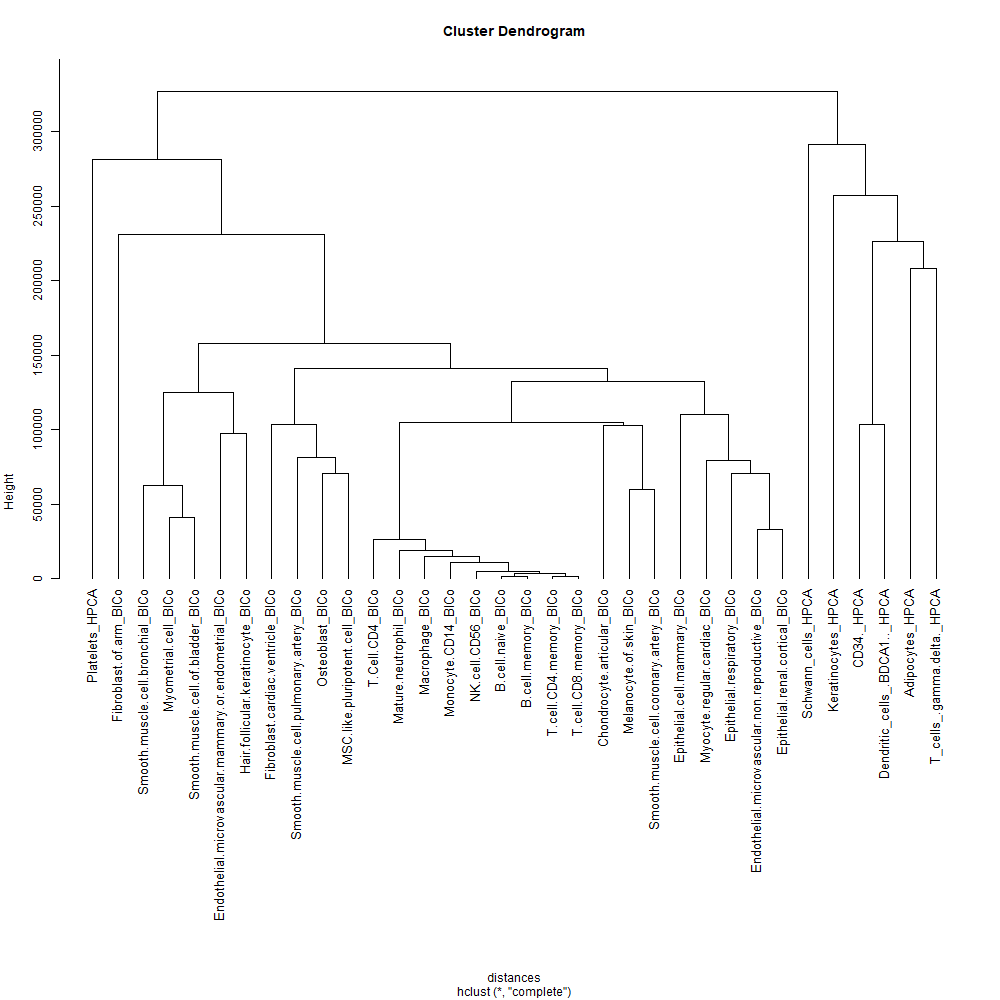

Supplementary Figure 7. Clusters formed when the BluePrint, ENCODE, and Human Primary Cell Atlas (HPCA) matrices are combined, quantile normalized, then clustered. HPCA cell types do not cluster with similar cell types in BluePrint or ENCODE, and instead form an external cluster of only HPCA profiles. This is likely due to batch or platform specific effects. We explored using this combined reference to produce predictions for the GTEx database (see Supplementary Figure 7), but instead took a different approach, as described in the main paper.

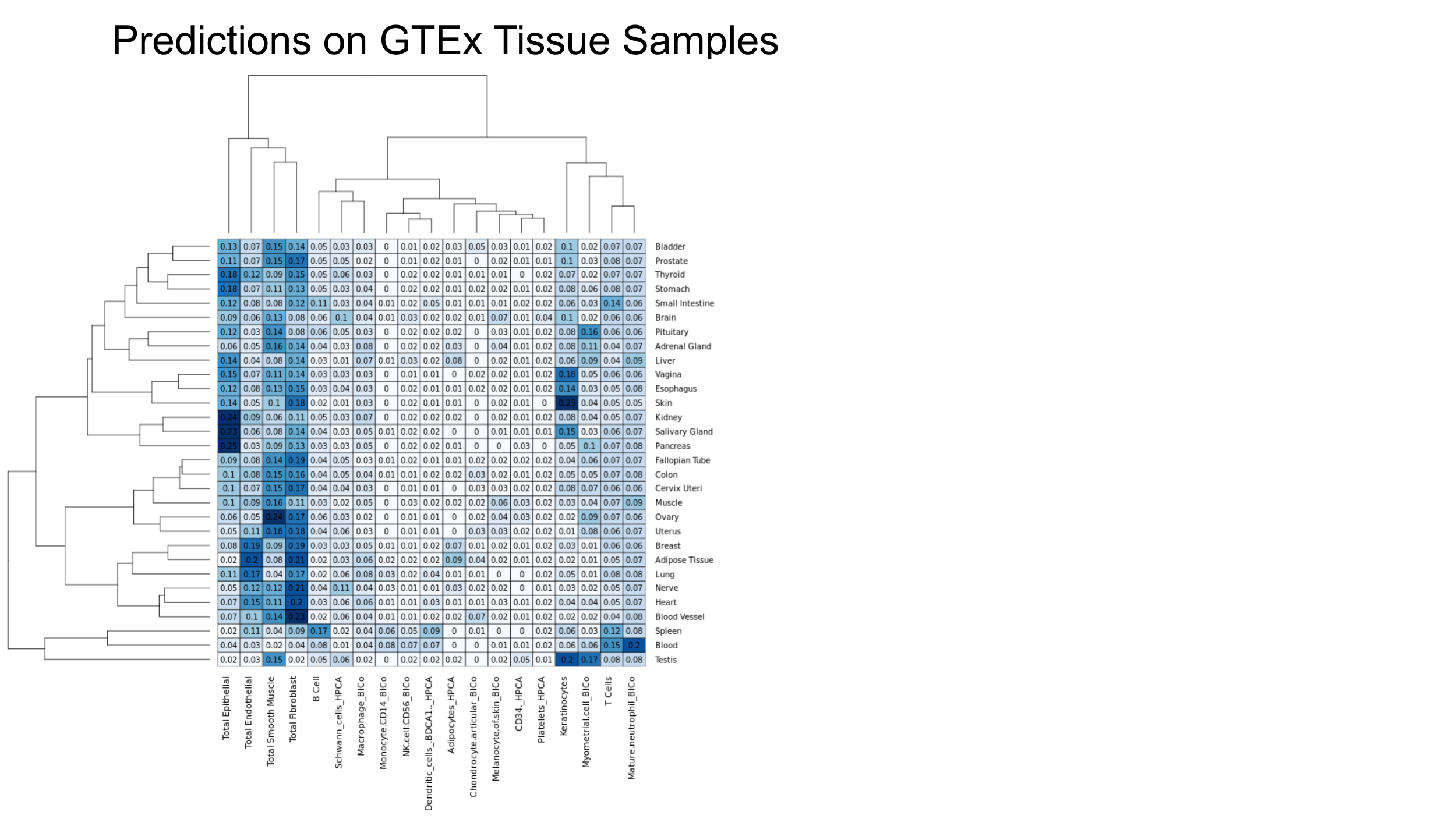

Supplementary Figure 8. Results of GEDIT when applied to the GTEx database when using a combined reference from the Human Primary Cell Atlas, BluePrint, and ENCODE. Six reference profiles from the HPCA were added to the BlueCode reference matrix, then all profiles were quantile normalized. GEDIT was run on the 17,382 samples from the GTEx database. Predicted cell type composition is averaged for all samples of the same tissue (right side of graph).
